# Modulation of RNA condensation by the DEAD-box protein eIF4A

**DOI:** 10.1101/689802

**Authors:** Devin Tauber, Gabriel Tauber, Anthony Khong, Briana Van Treeck, Jerry Pelletier, Roy Parker

**Author notes:** These authors contributed equally (Devin Tauber, Gabriel Tauber).

## Abstract

Stress granules are condensates of non-translating mRNAs and proteins involved in the stress response and neurodegenerative diseases. Stress granules form in part through intermolecular RNA-RNA interactions, although the process of RNA condensation is poorly understood. *In vitro*, we demonstrate that RNA is effectively recruited to the surfaces of RNA or RNP condensates. We demonstrate that the DEAD-box protein eIF4A reduces RNA condensation *in vitro* and limits stress granule formation in cells. This defines a purpose for eIF4A to limit intermolecular RNA-RNA interactions in cells, thereby allowing for proper RNP function. These results establish an important role for DEAD-box proteins as ATP-dependent RNA chaperones that can limit the intermolecular condensation and entanglement of RNA, analogous to the function of proteins like HSP70 in combatting protein aggregates.

**eTOC Blurb:** Stress granules are formed in part by the process of RNA condensation, which is mediated by and promotes *trans* RNA-RNA interactions. The essential DEAD-box protein and translation initiation factor eIF4A limits stress granule formation by reducing RNA condensation through its function as an ATP-dependent RNA binding protein, behaving analogously to how protein chaperones like HSP70 combat protein aggregates.

**Highlights:**

- RNA condensates promote intermolecular RNA-RNA interactions at their surfaces
- eIF4A limits the recruitment of RNAs to stress granules in cells
- eIF4A reduces the nucleation of stress granules in cells
- Recombinant eIF4A1 inhibits the condensation of RNA *in vitro* in an ATP-dependent manner

## INTRODUCTION

Eukaryotic cells contain ribonucleoprotein (RNP) granules in the nucleus and cytosol, including P-bodies (PBs) and stress granules (SGs) (Anderson and Kedersha, 2006; Banani et al., 2017). Stress granules are cytosolic RNP condensates composed of non-translating RNPs, which are involved in the stress response, neurodegeneration and viral infection (Protter and Parker, 2016; Ivanov et al., 2019). SGs typically form in response to translational shutoff induced by noxious stimuli such as arsenite, heat shock, and endogenous inflammatory molecules like prostaglandins, which all lead to phosphorylation of eIF2α and the activation of the integrated stress response (Aulas et al., 2017; Tauber and Parker, 2019). Stress granules can also form independently of eIF2α phosphorylation in response to inhibition of the eIF4F complex or high salt stress (Aulas et al., 2017). SGs and other RNP condensates are thought to form in part through multimeric RNA binding proteins crosslinking RNPs into larger networks (Banani et al., 2017; Shin and Brangwynne, 2018).

Recent evidence has emerged showing that intermolecular RNA-RNA interactions are involved in SG formation. SGs require a substantial pool of non-translating RNA to form (Protter and Parker, 2016). Thus, elevating the non-translating RNA concentration by injecting exogenous RNA induces SGs (Mahadevan et al., 2013). Furthermore, certain RNAs can seed foci in human cell lysates that recruit many SG proteins (Fay et al., 2017). Strikingly, modest concentrations of yeast total RNA readily condense in physiological salt and polyamine conditions, recapitulating the SG transcriptome in a protein-free context (Khong et al., 2017; Van Treeck et al., 2018), arguing that intermolecular RNA-RNA interactions contribute to SG formation. mRNPs are recruited to SGs in a biphasic manner, first engaging in transient docking interactions with the SG surface, which then transition into stable locking interactions that leave the RNA immobile within the granule (Moon et al., 2019), implying that surface recruitment of RNAs to SGs is a precursor to a more stable RNP assembly.

Roles for RNA in forming, maintaining, and organizing RNP condensates are not limited to SGs. For example, nuclear paraspeckles require the lncRNA *NEAT1* to form (Fox and Lamond, 2010; West et al., 2016). Moreover, specific RNA-RNA interactions between *NEAT1* domains may be important for paraspeckle formation and integrity (Lu et al., 2016; Lin et al., 2018). Similarly, the formation of RNA foci of repeat expansion RNAs has been suggested to occur through an RNA-driven condensation (Jain and Vale, 2017). In the *Drosophila* embryo, homotypic intermolecular base-pairing between *oskar* or *bicoid* mRNAs determines whether the RNAs will localize to RNP granules in the anterior or the posterior of the embryo, promoting polarity establishment (Ferrandon et al., 1997; Jambor et al., 2011). Specific intermolecular interactions between mRNAs may also be important for their recruitment to polarized RNP granules in *Ashbya gossypii* (Langdon et al., 2018). Despite their relevance to RNP condensate formation in cells, the properties of condensed RNA are not well-understood.

The diversity of intermolecular RNA-RNA interactions relevant to RNP granules and the low barriers for RNA condensation *in vitro* (Van Treeck and Parker, 2018) are consistent with RNA-based assemblies playing myriad cellular roles, but also create a need for the cell to regulate the processes of RNA condensation and recruitment. One possible modulatory mechanism would be the activity of RNA helicases, ATPases that can unwind RNA-RNA interactions and could thereby limit RNA condensation in the cell (Jarmoskaite and Russell, 2014). An interesting class of RNA helicases are members of the “DExD/H-box” family, which typically show highly cooperative binding to ATP and RNA but display low affinities for RNA when complexed with ADP, effectively making them ATP-dependent RNA binding proteins (Andreou and Klostermeier, 2013; Putnam and Jankowsky, 2013). Modulation of their ATPase activity can therefore control the rearrangement of RNPs (Hodge et al., 2011; Noble et al., 2011; Jankowsky et al., 2001; Putnam and Jankowsky, 2013). DEAD-box proteins typically disrupt structures in RNAs in part through ATP-dependent RNA binding, with ATP hydrolysis acting as a switch to release the protein from the RNA (Putnam and Jankowsky, 2013). Virtually all RNP granules contain DEAD-box proteins (Charroux et al., 1999; Saitoh et al., 2004; Dias et al., 2010; Hubstenberger et al., 2013; Calo et al., 2015; Tu and Barrientos, 2015; Jain et al., 2016; Hubstenberger et al., 2017; Markmiller et al., 2018; Youn et al., 2018), and these helicases are often conserved across eukaryotes (Linder and Fuller-Pace, 2013; Jarmoskaite and Russell, 2014). For example, SGs contain multiple conserved helicases, including eIF4A/Tif1 and DDX3/Ded1, in both mammals and yeast, respectively (Jain et al., 2016). Similarly, DEAD-box proteins are found localized in bacterial RNP granules as well (Al-Husini et al., 2018; Hondele et al., 2019).

Herein, we examine RNA condensation *in vitro* and SG formation in cells and how those processes are modulated by the essential translation initiation factor, SG component, and archetypal DEAD-box protein eIF4A. We show that RNA and RNP condensates stably interact with other RNAs or RNA-based condensates at their surfaces promoting RNA condensate assembly. In contrast, we demonstrate that eIF4A limits the recruitment of RNAs to SGs combating this process. Moreover, we show that the ATP-dependent binding of eIF4A to RNA, separate from its role in translation, can prevent the formation of SGs in cells, limit the interactions of PBs and SGs, and reduce the condensation of RNA in physiological conditions *in vitro*. Consistent with the mechanism of RNA unwinding by DEAD-box proteins, we show that RNA binding of eIF4A can be sufficient to limit RNA condensation or SG formation, but ATP hydrolysis may increase the effectiveness of eIF4A in limiting RNA condensation by allowing multiple cycles of RNA binding. Together, these data show that eIF4A functions as an ATP-dependent RNA binding protein to limit inappropriate intermolecular RNA-RNA interactions. Such an ATP-dependent RNA chaperone function is analogous to the role of protein chaperones, like HSP70, in limiting inappropriate intermolecular protein-protein interactions.

## RESULTS

To examine the nature of RNA condensation and how that process might affect RNP granule formation, we examined model RNA condensate systems *in vitro* with synthetic or purified RNAs. A wide variety of RNAs form RNA condensates *in vitro*, including all four homopolymers (Langdon et al., 2018; Van Treeck et al., 2018; Jain and Vale, 2017; Aumiller et al., 2016), demonstrating RNA condensation can robustly occur using non-Watson Crick interactions between bases. We examined the co-assembly of pairwise combinations of the homopolymer RNAs polyU, polyC, or polyA, which individually condense into RNA droplets of increasing viscosity, respectively (Van Treeck et al., 2018). The differential recruitment of short fluorescent oligonucleotides into homopolymer assemblies (Figure S1A & B) allowed us to distinguish between homopolymer condensates. The robustness of oligonucleotide recruitment coincided with the relative strengths of intermolecular RNA-RNA interactions between the oligonucleotides and the homopolymer scaffolds, with the most efficient recruitment occurring when the oligonucleotide could form Watson-Crick base pairs with the condensate scaffold. Some non-complementary RNAs were robustly recruited to condensate surfaces instead, such as oligoG with polyA or PTR with polyC (Figure S1A & B). These RNAs have some capacity for homotypic intermolecular interactions; for example, oligoG could in principle form intermolecular G-quadruplexes, suggesting that surface recruitment may occur when intermolecular RNA-RNA interactions between the client RNAs on the surface are preferred over interactions with the homopolymer scaffold or solvent.

Examination of the condensation of mixed homopolymers revealed that increasing the net RNA-RNA interaction strength through extensive base-pairing between polyA and polyU leads to gelatinous aggregates containing both homopolymers (Figure 1A). In contrast, mixtures of polyC and polyU spontaneously self-organize into patterned networks of alternating, immiscible polyC and polyU droplets (Figure 1A). A similar, but less organized, flocculation of polyA and polyC assemblies occurred (Figure 1A). These observations demonstrate that RNA-RNA interaction strengths determine the material phase of RNA condensates and that differential interaction strengths can lead to the homotypic clustering and specific compartmentalization of RNAs. Such sequence-specific organization of RNAs in condensates may be relevant to the homotypic clustering and/or organization of RNAs in RNP condensates in cells, such as the *Drosophila* germ granule (Trcek et al., 2015).

**Figure 1.**
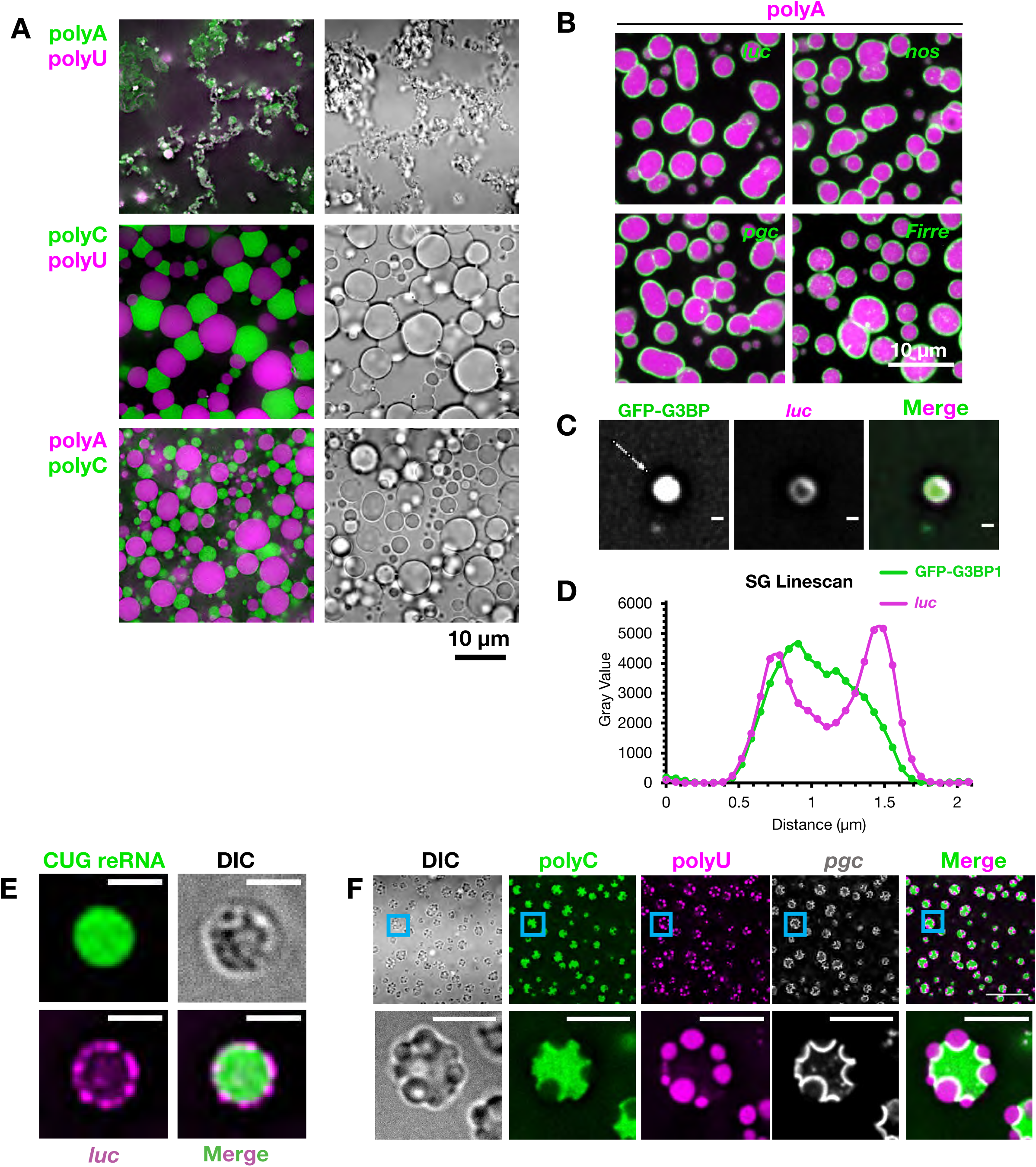
RNAs are recruited to and self-organize on RNA condensate surfaces. (A) Pairwise combinations of homopolymers were condensed together and visualized with fluorescent antisense oligos. (B) PolyA assemblies (labeled by U_19_-Cy3) were condensed with fluorescent *in vitro* transcribed RNAs. (C) Purified SG cores containing GFP-G3BP1 were incubated with fluorescent *luc* RNA. Scale bars are 500 nm. (D) Line profile of (C) along the line denoted by the white arrow. (E) A fluorescent myotonic dystrophy repeat RNA (reRNA) containing exons 5-11 of *DMPK* and ∼590 CUG repeats was condensed with fluorescent *luc*. Scale bars are 2 µm. (F) Fluorescent *pgc* was condensed with polyC and polyU (and the corresponding fluorescent antisense oligos), localizing to the polyC/polyU interface. Scale bars are 5 µm. *See also Figure S1*.

Multiple observations indicate that RNA condensates associate with one another at interfaces to minimize the total Gibbs free energy of their surfaces, which is the energetic cost of forming an interface between two phases (equivalent to the integral of the interfacial tensions with respect to area (Rowlinson and Widom, 1982)). First, the polyC/polyU interface is elongated, deforming the droplets from the typically favorable spherical shape, and forming regions with all three phases in contact (Figure 1A). This implies that the polyC/polyU interaction reduces the total Gibbs surface free energy of each individual droplet pair and the whole system in comparison to an equivalent system of dispersed droplets lacking the interfacial interactions (Rowlinson and Widom, 1982). A similar, but less extensive, docking of polyA and polyC condensates is also observed (Figure 1A), also consistent with interfacial interactions reducing the total surface free energy. In addition, time-lapse microscopy shows that polyC and polyU condensates are associated from an early time point and appear to nucleate off of each other (Movie S1), indicating that their interfacial association is not merely due to the crowding of droplets. Finally, polyC and polyU droplets remain associated with each other following mechanical disruption (Figure S1C), indicative of a bona fide physical interaction between the two condensate types.

The interactions of heterotypic RNA condensates raised the possibility that the recruitment of RNAs to RNA/RNP condensate surfaces would also occur. To test this possibility, we condensed polyA, polyC, or polyU in the presence of fluorescently labeled mRNAs. We observed that most mRNAs localized to the surfaces of all three homopolymer condensates, though some RNAs were internalized in polyC condensates (Figures 1B and S1D). Similarly, we observed that SGs isolated from mammalian cells (Figures 1C & D), and RNA assemblies formed from a myotonic dystrophy associated CUG repeat RNA (known to form RNA condensates in cells; Jain and Vale, 2017), recruit RNA to their surfaces *in vitro* (Figure 1E). Thus, a diversity of RNA and RNP assemblies recruit RNAs to their surfaces.

We also observed that the condensation of *pgc* RNA with both polyC and polyU led to the formation of polyC/polyU multi-phase assemblies (Figure 1F) with *pgc* RNA localized robustly to the polyC/polyU interface. This observation argues that specific RNAs might act similarly to surfactants or interfacial shells between distinct RNA/RNP condensate phases, and thereby stabilize multi-phase RNP condensates like the nucleolus (Feric et al., 2016).

Three observations demonstrate that the recruitment of RNAs to the surface of an RNA condensate leads to enhanced interaction between those RNAs and the formation of an RNA shell of enhanced stability enveloping the underlying RNA condensate. First, the surface ring of *pgc* mRNAs recruited to polyA condensates is more stable following dilution than the underlying polyA assembly, persisting for several minutes longer (Figures 2A & B, Movie S2). Second, while the internal polyA droplet (visualized by fluorescent oligoU) is somewhat dynamic by FRAP analysis (mobile fraction at 15 min post-photobleaching, m.f. = 0.29 ± 0.02), the *pgc* shell shows virtually no recovery (m.f. = 0.02 ± 0.02), implying that the shell is immobile, which we confirmed by photobleaching only half the condensate and observing no *pgc* diffusion (Figure S2). Third, using 4’-aminomethyltrioxsalen, which crosslinks RNA duplexes in UV light (Frederikson and Hearst, 1979), we observed that intermolecular *pgc*-*pgc* crosslinking, analyzed on denaturing gels, increased when assembled on the surface of polyA condensates as compared to in dilute solution, or even when compared to homotypic, self-assembled *pgc* gels (Figures 2C-E).

**Figure 2.**
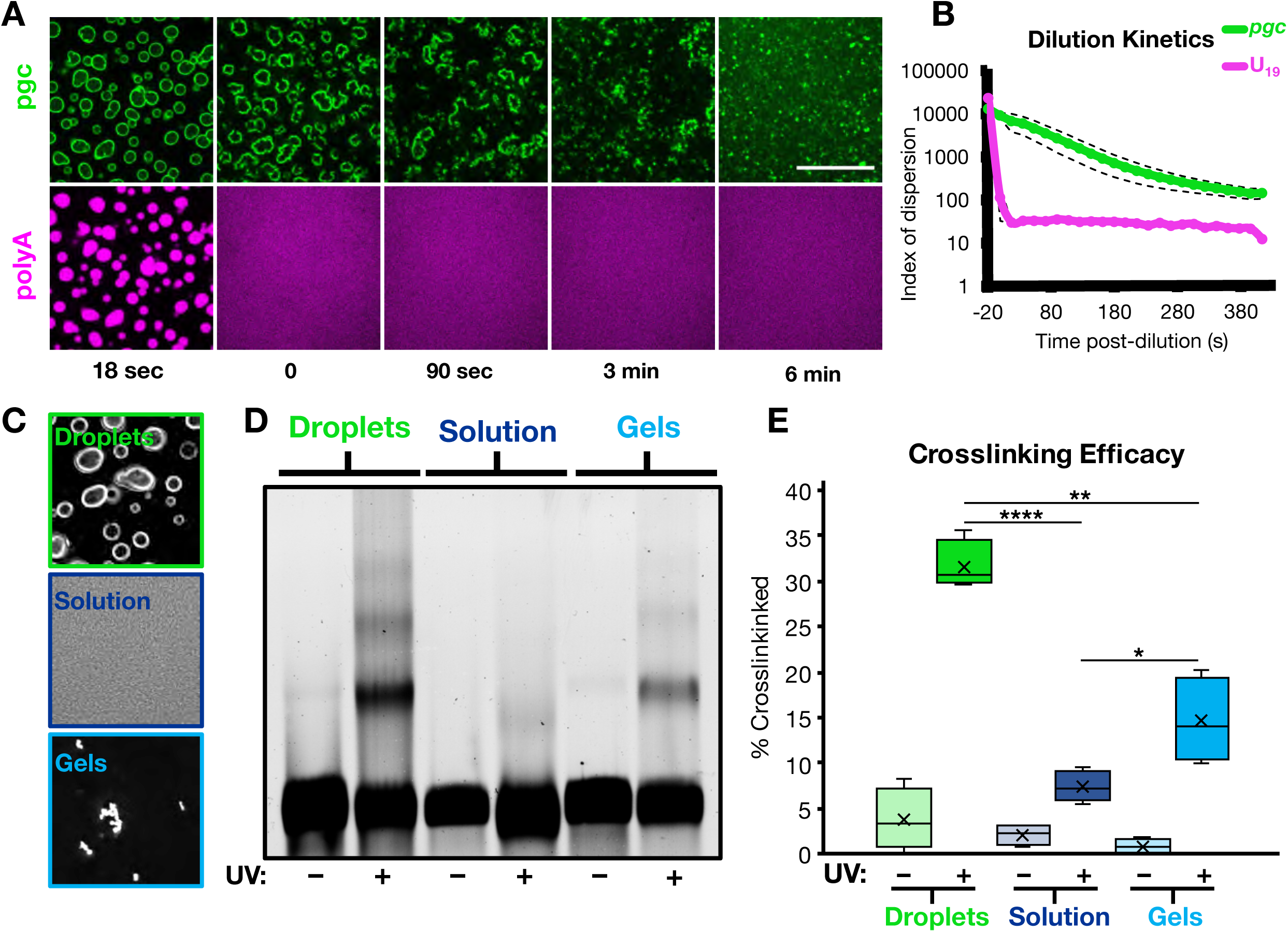
RNA condensate surface localization stabilizes intermolecular RNA-RNA interactions. (A) Fluorescent *pgc* was condensed with polyA and fluorescent U_19_, and the condensates were subjected to 1:10 dilution in TE buffer. Scale bar is 20 µm. (B) Quantification of (A) as an index of dispersion, showing the persistence of *pgc* shell assemblies over time. Dashed lines are 95% confidence intervals. *n* = 6 replicates. (C) Images of the crosslinking conditions showing *pgc* on droplets, in solution, and as gels. (D) Representative fluorescence denaturing gel. (E) Quantification of (D), showing that RNA condensate surfaces enhance intermolecular RNA interactions in comparison to solvated RNA or RNA condensed alone. X represents the mean. \**p*<0.05, \*\**p*<0.01, \*\*\*\**p*<10^−4^. *n =* 4 replicates. *See also Figure S2*.

These observations argue that the surfaces of RNA assemblies recruit RNAs. Moreover, the binding of RNAs onto the surface of an RNP granule can concentrate RNAs/RNPs, thereby promoting additional intermolecular RNA/RNP interactions, further stabilizing the RNA/RNP condensate. This raises the possibility that recruitment of RNAs to the surfaces of RNP granules in cells will be a robust process, and therefore mechanisms must exist to limit the recruitment of RNAs onto the surfaces of RNP granules. We hypothesized that one or more DEAD-box proteins would act as ATP-dependent RNA chaperones in cells by limiting or dissociating the intermolecular RNA-RNA interactions that promote RNP condensation.

### eIF4A limits RNP partitioning into stress granules

To examine the role of DEAD-box proteins functioning as RNA chaperones in limiting RNA condensation, we focused on SGs. Multiple DEAD-box proteins partition into SGs (Chalupníková et al., 2008; Hillicker et al., 2011; Jain et al., 2016; Markmiller et al., 2018) with eIF4A1, an essential, highly-conserved component of the eIF4F translation initiation complex, being the most abundant in both U-2 OS and HeLa cell lines (Figure 3A; Beck et al., 2011; Itzhak et al., 2016). We hypothesized that eIF4A would have additional functions since eIF4A1 is present at ∼10X the concentration of other eIF4F components (Itzhak et al., 2016; Beck et al., 2011; Pause et al., 1994), with between 5-50 molecules of eIF4A1 for each mRNA in U-2 OS and HeLa cells (calculations in STAR Methods).

**Figure 3.**
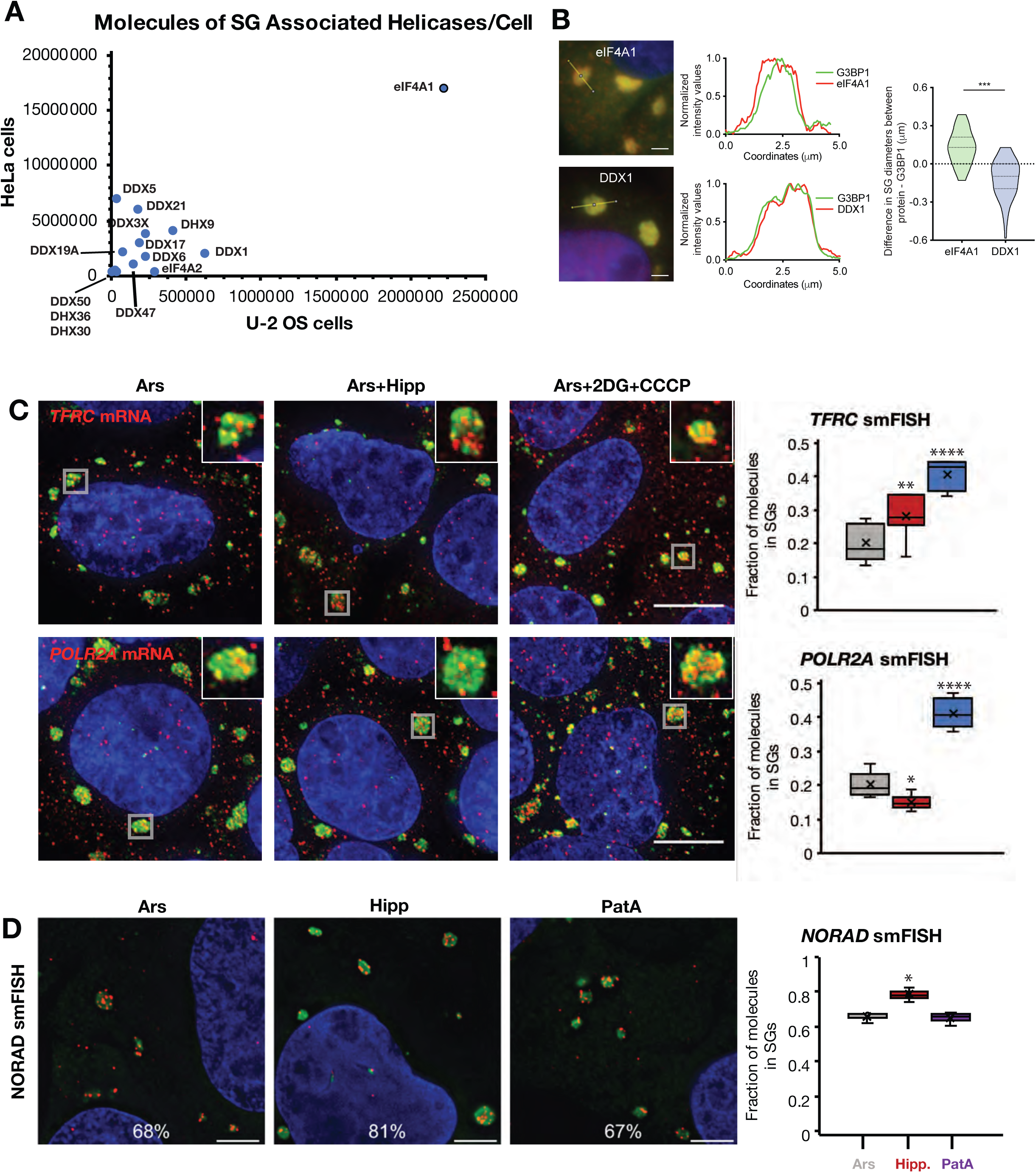
eIF4A limits RNA recruitment to SGs. (A) Scatterplot of SG-associated helicase abundance in U-2 OS vs. HeLa cells. (B) Immunolocalization of eIF4A1 in SGs. eIF4A1 extends past the SG periphery as compared to DDX1, which co-extends with G3BP1. Scale bars are 2 µm. *n* = 3 replicates. (C) smFISH images and quantification of SG enrichment for *TFRC* and *POLR2A* mRNAs in U-2 OS cells treated with 60’ arsenite, then 30’ DMSO, hippuristanol, or 2DG and CCCP. Gray boxes denote the SGs in the insets. SGs are visualized by anti-PABPC1 IF. Scale bars are 20 µm. *n* ≥ 5 frames. (D) *NORAD* lncRNA smFISH images and quantification in U-2 OS cells treated with arsenite, hippuristanol, or PatA. SGs are visualized by anti-G3BP1 IF. *n* = 3 replicates. \**p*<0.05, \*\**p*<0.01, \*\*\**p<*10^−3^, \*\*\*\**p*<10^−4^. *See also Figure S3*.

eIF4A1 partitions into SGs during arsenite stress, and on average extends further into the cytosol then the traditional SG marker, G3BP1 (Figure 3B), indicating eIF4A1 is enriched at the periphery of SGs. In contrast, DDX1, another SG RNA helicase, is uniformly distributed within the granule and overlaps with G3BP1 (Figure 3B). The peripheral concentration of eIF4A1 in SGs is consistent with eIF4A1 modulating the interactions of RNAs with the surfaces of SGs.

If eIF4A limits the condensation of RNAs into SGs, then eIF4A inhibition should increase RNA partitioning into SGs. To separate the effects of eIF4A inhibition on translation initiation from its helicase function, we first treated U-2 OS cells with arsenite to inhibit bulk translation initiation through eIF2*α* phosphorylation, then added hippuristanol for 30’, which specifically inhibits the helicase activity of eIF4A by preventing ATP-dependent RNA binding (Bordeleau et al., 2006a; Lindqvist et al., 2008). We then examined the partitioning of mRNAs into SGs by single molecule fluorescence in situ hybridization (smFISH).

We observed that the fraction of *TFRC*, but not *POLR2A*, mRNAs associated with SGs increased after hippuristanol treatment (Figure 3C). We also observed that bulk mRNAs (as judged by oligo(dT) FISH) and *DYNH1C1*, *AHNAK*, and *PEG3* mRNAs, but not the MCM2 mRNA, partition more strongly into SG in the presence of arsenite and hippuristanol as compared to arsenite alone (Figure S3B). Since there are additional helicases in SGs, we also examined mRNA partitioning into arsenite-induced SGs following ATP depletion, which would inhibit all ATP-dependent RNA helicases. After ATP depletion, we observed both *TFRC* and *POLR2A* mRNAs increased their partitioning into SGs, consistent with other ATP-dependent mechanisms, including additional RNA helicases, limiting mRNA condensation into SGs (Figure 3C). We note the caveat that ATP depletion affects many cellular processes, and hence there may be indirect effects from ATP depletion. However, given the effects of other DEAD-box proteins on RNP granules (see below), it is highly probable that the increase in RNA partitioning we observe is due in part to inhibition of other ATP-dependent RNA chaperones.

To examine the role of eIF4A helicase activity on RNA partitioning into SGs independently of its role in translation in a second context, we also measured the partitioning of the SG-enriched (Khong et al., 2017) *NORAD* lncRNA, which is not thought to be translated, in the presence of arsenite, hippuristanol, or Pateamine A (PatA), which inhibits eIF4A’s function in translation while stimulating eIF4A RNA binding and helicase activity (Bordeleau et al., 2005; Bordeleau et al., 2006b). We observed that *NORAD* RNA increased partitioning to SGs during hippuristanol treatment, as compared to arsenite or PatA (Figure 3D). Similarly, we observed bulk mRNA and the abundant RNA binding protein G3BP1 partitioned more robustly to SGs in hippuristanol-treated cells as assessed by oligo(dT) FISH or IF, respectively, while average SG sizes where slightly decreased with hippuristanol or pateamine A (Figure S3B, C). This suggests that the density of SGs is increased with inhibition of eIF4A.

These observations demonstrate that the RNA helicase activity of eIF4A1, independent of its role in translation, limits the accumulation of RNPs in SGs. We suggest that RNAs not affected by eIF4A such as POLR2A or MCM2 might be limited from entering stress granules by other RNA chaperones or helicases or might be primarily targeted to SGs by proteins. For example, DEAD-box proteins can preferentially bind, target, and/or unwind differently structured RNAs *in vitro* and *in vivo* (Murat et al., 2018; Ribero de Almeida et al., 2018; Chen et al., 2018), and these preferences could extend to the ability of DEAD-box proteins to limit the condensation of specific RNAs.

### eIF4A limits stress granule formation

We hypothesized that limiting intermolecular RNA interactions through eIF4A would affect not only the recruitment of mRNPs to SGs, but also intermolecular RNA-RNA interactions between mRNPs that contribute to SG assembly. To test this possibility, we took advantage of the observation that during arsenite stress, SGs do not form in cells lacking the abundant RNA binding proteins G3BP1/G3BP2, which provide protein-protein interactions to assist SG assembly through G3BP dimerization (Tourrière et al., 2003; Kedersha et al., 2016). We hypothesized that inhibition of eIF4A would compensate for the absence of G3BP in SG formation by promoting increased levels of intermolecular RNA-RNA interactions. Thus, we examined SG formation in WT and ΔΔG3BP1/2 cell lines during arsenite stress, where we altered eIF4A function after 30’ of arsenite stress with either hippuristanol, blocking RNA binding by eIF4A, or PatA as a negative control. Puromycin labeling revealed that translation was repressed similarly in all three conditions, indicating that any effects are not due to additional translational repression (Figure S4A).

Importantly, hippuristanol treatment partially restored SG formation in the ΔΔG3BP1/2 cell lines as assessed by PABPC1 IF, but PatA treatment did not (Figure 4A, B). These PABPC1 foci are SGs since they are sensitive to cycloheximide (Figure S5A), which blocks SG formation by trapping mRNAs in polysomes (Kedersha et al., 2000), they do not colocalize with PBs, and they contain multiple SG proteins, polyadenylated RNA, and the SG-enriched RNAs *AHNAK* and *NORAD* (Figure S5B-C). Depletion by siRNAs of eIF4A1 (Figure S4B), the predominant form of eIF4A in U-2 OS cells (Figure 3A), also led to the restoration of SGs in the ΔΔG3BP1/2 cell line during arsenite treatment (Figure 4B, C). This demonstrates that eIF4A limits SG formation separately from promoting translation.

**Figure 4.**
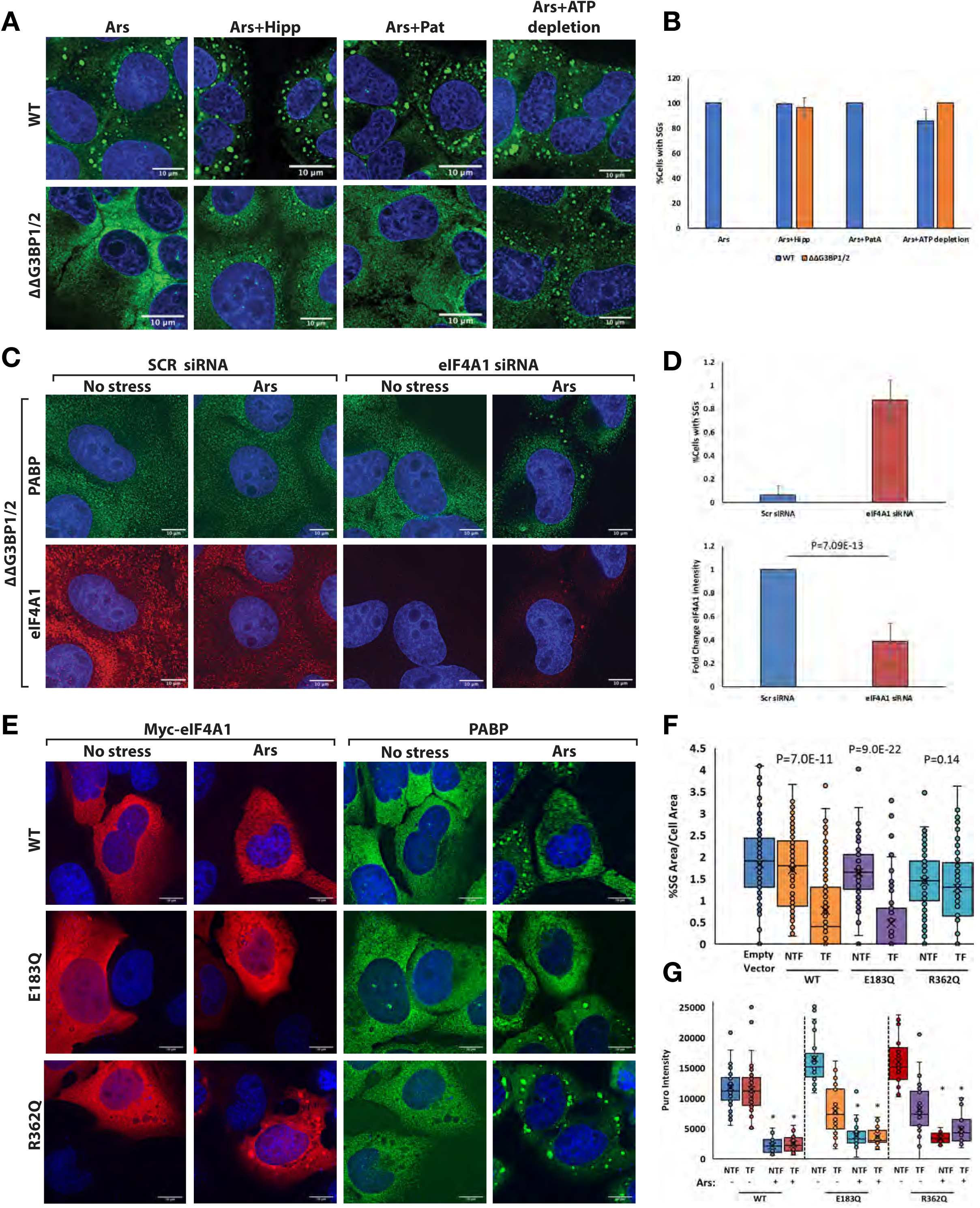
eIF4A limits SG formation. (A) Images displaying SG formation (as assessed by PABPC1 IF) in both wildtype and ΔΔG3BP1/2 U-2 OS cells. Arsenite, alone or in combination with hippuristanol or PatA robustly induces SG formation in wild-type cells. Hippuristanol, following arsenite, can induce SG formation in ΔΔG3BP1/2, indicating that inhibition of eIF4A’s helicase activity can partially restore SG formation independently of translational shutoff. Additionally, arsenite followed by ATP depletion restores SGs in ΔΔG3BP1/2 cells to a greater extent than arsenite and hippuristanol, suggesting other ATP driven machines limit RNA condensation. (B) Quantification of images in (A). (C) siRNA knockdown of eIF4A1 (quantified by fold change in eIF4A1 intensity in (D)) in ΔΔG3BP1/2 cells restores SGs upon addition of arsenite, indicating that eIF4A1 limits SG formation. (E) Overexpression of eIF4A1 in wildtype U-2 OS cells prevents SG formation, as quantified by %SG area/cell area in cells expressing Myc-tagged eIF4A1, compared to non-transfected neighbor cells or control transfections. ATPase mutant E183Q was able to prevent SG formation when over-expressed, however, RNA-binding mutant R362Q was not. (F) Quantifications of (E) represented by %SG Area/Cell Area of Myc-eIF4A1 over-expressing (TF) or non-transfected (NTF) cells. (G) Quantifications of translation between NTF and TF WT and mutant versions of Myc-eIF4A1 as assessed by puromycin intensity. Notice that translation is repressed equally in all conditions when arsenite is added, indicating that effects on SG formation are not due to increased translation (\**p* ≤ 0.05). For each experiment, error bars represent standard deviations, with *n* ≥ 3 replicates. *See also Figure S4 and S5*.

In principle, eIF4A could limit SG formation by limiting intermolecular RNA-RNA interactions, or by increasing the off-rate of RNA-binding proteins contributing to SG assembly. However, inhibition of eIF4A helicase activity with hippuristanol does not alter the exchange rates of G3BP1 (Figure S4H), consistent with eIF4A altering SG formation by promoting the dissociation of intermolecular RNA-RNA interactions.

Since SGs were only partially restored with hippuristanol, we also depleted cells of ATP after 30’ of arsenite stress, when ATP is no longer necessary to release mRNAs from polysomes (Jain et al., 2016; Khong and Parker, 2018), to inhibit all ATP-dependent DEAD-box proteins. Depletion of ATP restored SG formation more robustly than hippuristanol in ΔΔG3BP1/2 cells (Figure 4A). We interpret these observations to signify that in addition to eIF4A, other ATP-dependent mechanisms limit SG formation, although we cannot rule out indirect effects of ATP depletion on cell physiology that also lead to enhanced SG formation. The observation that ATP depletion following translational arrest promotes SG formation is consistent with RNP condensation into SGs being an energetically favored process.

Since inhibiting eIF4A increased SG formation independent of its role in translation, we predicted that over-expression of eIF4A would limit intermolecular RNA-RNA interactions and thereby limit SG formation. To test this, we over-expressed Myc-tagged eIF4A1 in wildtype U-2 OS cells by transient transfection (Figure S4C). We then examined whether cells with over-expressed eIF4A1 could still form SGs in response to arsenite treatment. We observed that cells with over-expression of eIF4A1 demonstrated defects in SG formation, although those cells still robustly repressed translation (Figure 4E, F, G, S4D), consistent with eIF4A1 limiting RNA condensation into SGs.

In principle, eIF4A could limit RNA condensation in two related manners. First, when bound to ATP it could bind RNA and thereby compete for RNA-RNA interactions. Alternatively, it could utilize the energy of ATP hydrolysis to limit RNA condensation. Since eIF4A is known to promote RNA duplex unwinding by binding RNA in an ATP-dependent manner (Rogers et al., 1999; Rogers et al., 2001), these are related mechanisms. In order to determine the mechanism by which eIF4A limits RNP condensation, we examined how mutations that inhibit either RNA binding or ATP hydrolysis affected the ability of eIF4A to limit SG formation when over-expressed in U-2 OS cells.

We observed that overexpression of an RNA binding mutant of eIF4A (R362Q [Pause et al., 1994]) did not prevent SG formation (Figure 4E-F). This indicates that RNA binding of eIF4A is required for the inhibition of SG formation. In contrast, we observed that overexpression of an ATPase inactive mutant (E183Q) that can still bind ATP and RNA (Pause et al., 1994; Svitkin et al., 2001; Oguro et al., 2003) repressed SG formation similarly to WT eIF4A (Figure 4E-F). This indicates that, at least when over-expressed, eIF4A does not require ATP hydrolysis to limit stress granule formation, suggesting a model whereby eIF4A is acting as an ATP-dependent RNA binding protein to inhibit SG formation, with ATP hydrolysis serving to release the RNA from eIF4A. These effects are independent of translation repression since cells with any of the eIF4A variants over-expressed repressed translation equally in response to arsenite treatment (Figure 4G).

Since other RNA helicases likely function to limit RNA condensation, we tested if a closely related DEAD-box protein, DDX19A/DBP5 (Cencic and Pelletier, 2016), could prevent SG formation in an analogous manner to eIF4A1. DDX19A over-expression has been reported to inhibit the formation of SGs induced by the transcriptional inhibitor tubercidin by an unknown mechanism (Hochberg-Laufer et al., 2019). We reasoned that DDX19A could also limit RNA condensation since its helicase core sequence is highly similar to eIF4A. Overexpression of mCherry-DDX19A inhibited arsenite-induced SG formation to a similar extent as eIF4A, while over-expression of mCherry alone did not decrease SG formation (Figure S4E, G). Similar to eIF4A, overexpression of RNA binding deficient DDX19A fails to inhibit SG formation, while overexpression of an ATP hydrolysis mutant still inhibits SG formation (Figure S4E, G). This demonstrates that multiple different ATP-dependent RNA chaperones can limit RNA condensation and SG formation, which is consistent with ATP depletion having a larger effect on SG formation than inhibition of eIF4A alone (Figure 4A).

### eIF4A limits docking of P-bodies with stress granules

Heterotypic RNP granules, like SGs and PBs (Kedersha et al., 2005) or nuclear speckles and paraspeckles (Fox et al., 2002), are observed to dock in cells. Since heterotypic RNA condensates minimize their surface free energy through RNA-RNA docking interactions (Figure 1A), we hypothesized that PB/SG interfaces might occur through intermolecular RNA interactions. If so, the docking of PBs and SGs would be predicted to be modulated by eIF4A. To test this possibility, we treated cells with arsenite, either alone or with hippuristanol or PatA treatment, and then quantified PB/SG interfaces. We normalized the docking frequency of PBs and SGs to either total numbers of PBs or SGs, or to total PB or SG area to control for the possibility that drug additions affected the amount or sizes of either RNP granule. We observed that the docking frequency of PBs and SGs increased with hippuristanol addition compared to arsenite or arsenite and pateamine A regardless of normalization (Figure 5), although when normalized to PB area the difference between hippuristanol and pateamine A was not statistically significant, probably because pateamine A led to a decrease in overall PB area (Figure 5F). These observations suggest that eIF4A can limit docking interactions between PBs and SGs.

**Figure 5.**
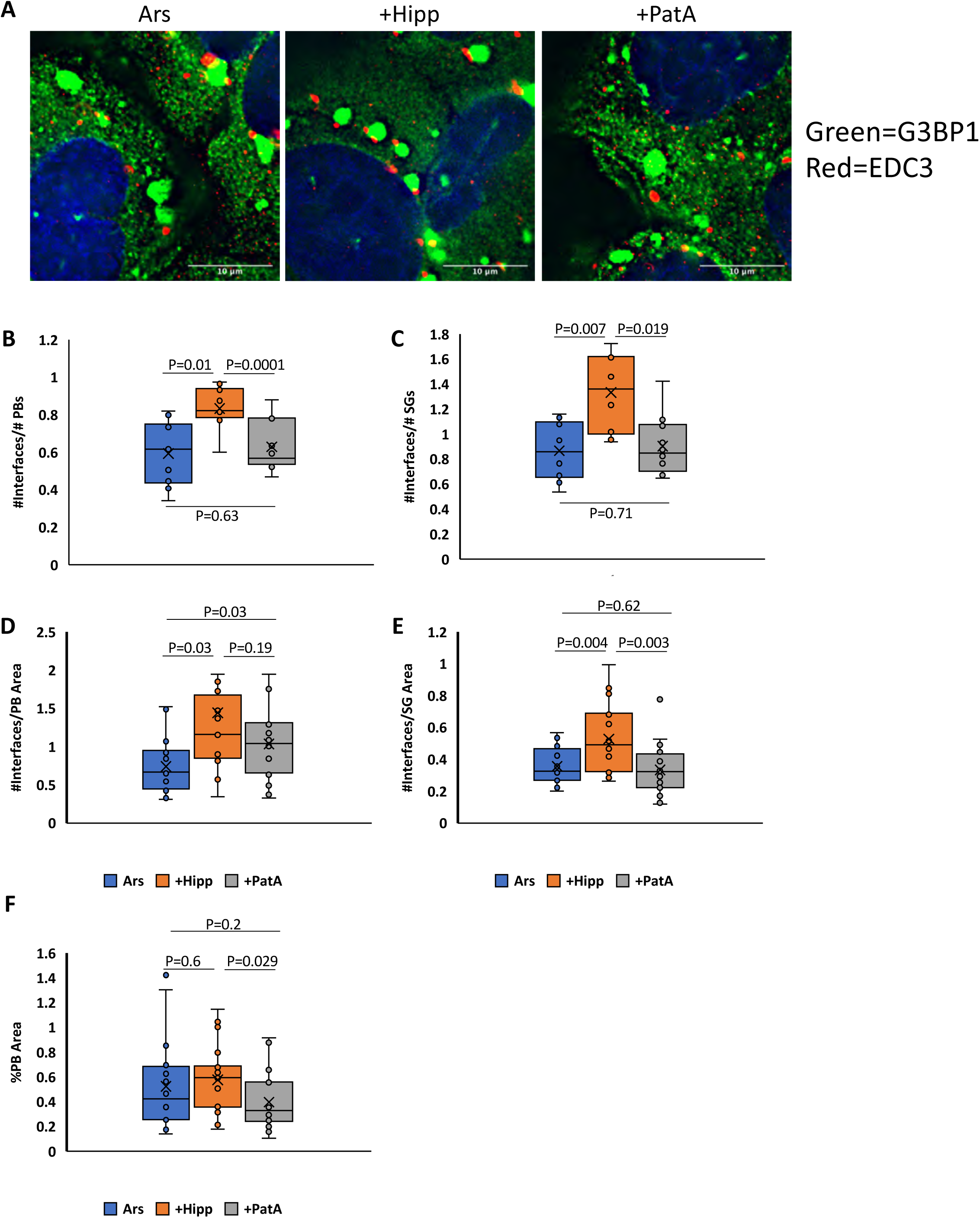
eIF4A limits the docking of P bodies with stress granules. (A) The number of PB/SG interfaces increase in the presence of arsenite in combination with hippuristanol compared to combinations of arsenite with PatA or arsenite alone. This effect holds true when interface quantities are normalized to the total amount or area of PBs (B, D) or SGs (C, E). PBs are visualized by anti-EDC3 IF and SGs by anti-G3BP1 IF. *n* = 5 replicates. Pateamine A addition produces a slight decrease in PB area (F).

### eIF4A1 and ATP are sufficient to limit RNA condensation *in vitro*

To determine if eIF4A was sufficient to limit RNA condensate formation *in vitro*, we examined how recombinant human eIF4A1 affected the formation of RNA condensates *in vitro*. We assessed RNA condensate formation by observing RNA condensates labelled with SYTO^TM^ RNASelect^TM^, an RNA specific fluorescent dye. RNA condensates were formed from total yeast RNA in physiological K^+^ and polyamine concentrations in the presence of PEG to simulate the crowded environment of the cell (Zimmerman, 1993; Ellis, 2001; Delarue et al., 2018). We performed these experiments with eIF4A at 10 μM, which is its approximate concentration in the cytosol (Itzhak et al., 2016) and with RNA at 150 μg/ml, which we have estimated is the concentration of exposed ORF RNA during acute translational inhibition (Van Treeck et al., 2018). We then examined the formation of RNA condensates over time in the presence or absence of recombinant eIF4A1, with ATP (1 mM) or the non-hydrolyzable ATP analog adenylyl-imidodiphosphate (ADPNP, 1 mM). We observed that ATP had a small effect on RNA condensation, perhaps because it is essentially a monovalent RNA and could compete for intermolecular RNA-RNA interactions (Figure S6).

An important result was that RNA condensation, as assessed by fluorescent labelling was strongly reduced by the addition of eIF4A1 and ATP as compared to ATP alone (Figure 6A-B). eIF4A1 in the absence of ATP had no effect on RNA condensation (Figure S6). This demonstrates that eIF4A1 in the presence of ATP can limit RNA condensation. The substitution of the non-hydrolyzable ADPNP for ATP still allowed eIF4A1 to limit RNA condensation, but less effectively (Figure 6A-B). Since eIF4A retains its ability to bind RNA when bound to ADPNP (Lorsch and Herschlag, 1998a), this is consistent with ATP-dependent RNA binding being sufficient to limit RNA condensation, and that process being increased in efficacy by ATP hydrolysis allowing eIF4A to perform multiple rounds of RNA binding. In contrast, the addition of ATP or ADPNP alone had small but similar effects on RNA condensation in the absence of eIF4A1 (Figure S6).

**Figure 6.**
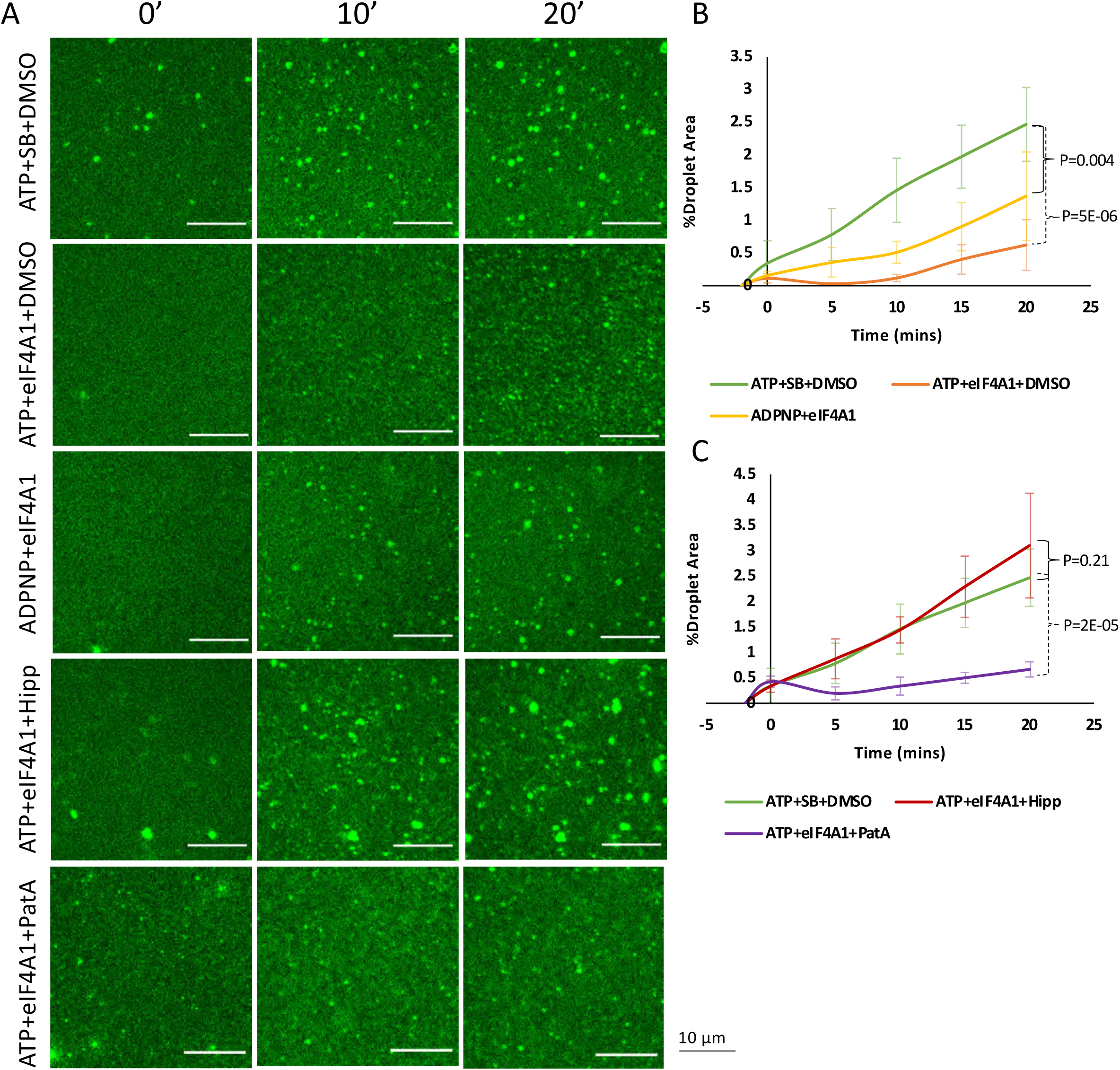
Recombinant eIF4A is sufficient to limit RNA condensation *in vitro*. (A) Formation of *in vitro* fluorescent total RNA droplets was monitored over a period of 20 minutes and is inhibited by eIF4A+ADPNP, and more drastically by eIF4A+ATP, suggesting that binding of eIF4A to RNA can prevent RNA condensation, with eIF4A catalytic activity being the most effective at inhibiting spontaneous RNA assembly. Hippuristanol can inhibit eIF4A1 catalytic function and restores droplet formation, while pateamine A does not. (B) quantification of RNA condensation kinetics comparing ATP + eIF4A1 and ADPNP + eIF4A1 and (C) hippuristanol addition compared to pateamine A as assessed by the percent of total area occupied by droplets (%Droplet area) per time. *n* = 3 replicates. SB = protein storage buffer. Error bars represent standard deviations of the mean. *For control images, see Figure S6*.

To further address the mechanism of RNA condensation inhibition by eIF4A *in vitro* we tested how hippuristanol and pateamine A affected eIF4A1 decondensase function. Without eIF4A1, hippuristanol (10 µM) or pateamine A (1 µM) in the presence of ATP had no effects on condensate formation (Figure S6). However, eIF4A1 preincubated with hippuristanol before RNA addition did not prevent condensate formation (Figure 6A, C). Since hippuristanol prevents eIF4A from binding RNA, this argues ATP and RNA binding are required for eIF4A to limit RNA condensation *in vitro*. In contrast, preincubation with pateamine A, which slightly increases the catalytic efficiency of eIF4A1 (Low et al., 2005), did not limit eIF4A1 function in preventing RNA condensate formation (Figure 6A, C). These data confirm our *in vivo* observations that hippuristanol, but not pateamine A, addition following translational inhibition in G3BP null cells can rescue SG formation by preventing eIF4A’s function in limiting RNA condensation.

## DISCUSSION

Several observations argue that RNA condensation into RNP granules is a thermodynamically favored process that is countered by energy-consuming processes in the cell. First, multiple distinct RNAs are robust at self-assembly *in vitro*, including under physiological conditions of salts and polyamines (Figure 1; Figure S1; Aumiller et al., 2016; Jain and Vale, 2017; Langdon et al., 2018; Van Treeck et al., 2018). Moreover, RNA generally requires lower concentrations to condense *in vitro* than intrinsically disordered proteins (Van Treeck and Parker, 2018). Second, we observe that condensates of different RNA composition can interact to reduce the surface free energy of the RNA condensates (Figure 1). Third, multiple RNA condensates or purified stress granules recruit RNAs to their surface *in vitro* (Figure 2). Furthermore, in cells, depletion of ATP promotes the condensation of untranslated mRNPs into SGs (Figure 3C), even in the absence of protein factors normally required for SG formation (Figure 4A), demonstrating cells utilize energy to limit SG formation.

*In vitro*, we observe that the recruitment of RNAs to the surface of an RNA condensate creates a high local concentration, which promotes the formation of additional interactions between the molecules to further stabilize the RNA assembly (Figure 2). This creates a positive feedback loop in RNA condensation. Thus, RNA condensation has the potential to create an “RNA entanglement catastrophe” wherein extensive RNA condensation would limit the proper distribution and function of RNAs. An implication of these observations is that cells must contain mechanisms to limit the intermolecular interactions and condensation of RNA.

We present several lines of evidence arguing that eIF4A functions to limit cytosolic RNA condensation. First, inhibition of eIF4A1 function by knockdown or hippuristanol treatment, but not PatA, restores SG formation in cells lacking G3BP without changes in translational repression (Figure 4A & B). Second, overexpression of eIF4A1 limits arsenite-induced SG formation, although translation remains repressed (Figure 4E-G, S4D). Third, inhibition of the RNA binding and ATPase activity of eIF4A by hippuristanol treatment during arsenite stress enriches bulk, and specific, RNA recruitment to stress granules (Figures 3 and S3). Fourth, inhibition of eIF4A RNA binding and ATPase activity with hippuristanol increases PB/SG docking events (Figure 5). Finally, recombinant eIF4A1 is able to limit RNA condensation *in vitro* in an ATP-dependent manner (Figure 6). Since inhibition of eIF4A helicase activity with hippuristanol does not alter the exchange rates of G3BP1 (Figure S4H), we interpret these observations to suggest that eIF4A limits RNA condensation by the disruption of intermolecular RNA-RNA interactions. Since RNA binding, but not ATP hydrolysis, is required for limiting SG formation when eIF4A1 is over-expressed (Figure 4E-F), RNA binding is the critical feature of eIF4A’s ability to limit RNA condensation.

These results synergize well with biochemical data on the eIF4A helicase mechanism. RNA duplex unwinding by eIF4A and other DEAD-box proteins is coupled not so much to ATP hydrolysis as to the ATP-dependent binding of eIF4A to the RNA substrate, which kinks RNA and displaces several nucleotides, locally destabilizing the duplex, which may then dissociate thermally (Rogers et al., 1999; Rogers et al., 2001; Liu et al., 2008; Putnam and Jankowsky, 2013). Hence, productive unwinding events are determined in part by the relative thermostability of the RNA•eIF4A•ATP complex compared to the structured RNA, limiting eIF4A’s ability to efficiently resolve duplexes more stable than ∼10-15 bp (Δ*G* ∼ –25 kcal/mol; Rogers et al., 1999; Rogers et al., 2001). This may provide insight into the nature of the *trans* RNA interactions that drive RNA condensation and recruitment to condensates, namely that they are individually rather weak, despite their apparent thermostability in summation. Thus, ATP serves as a molecular switch to increase the affinity of the protein for RNA, while ATP hydrolysis and P_i_ release cause conformational changes to lower the affinity of eIF4A for RNA, effectively making eIF4A function as an ATP-dependent RNA binding protein (Lorsch and Herschlag, 1998a; Lorsch and Herschlag, 1998b; Sun et al., 2014). Taken together, we suggest that while the binding of eIF4A to RNA is the key step in limiting RNA condensation, the ATPase-driven cycling of eIF4A on and off transcripts increases its ability to limit *trans* interactions by allowing multiple RNA binding events (Figure 7A). Accordingly, eIF4A limits RNA condensation as an ATP-dependent RNA binding protein, providing the cell with a controllable means of buffering and regulating intermolecular RNA interactions.

**Figure 7.**
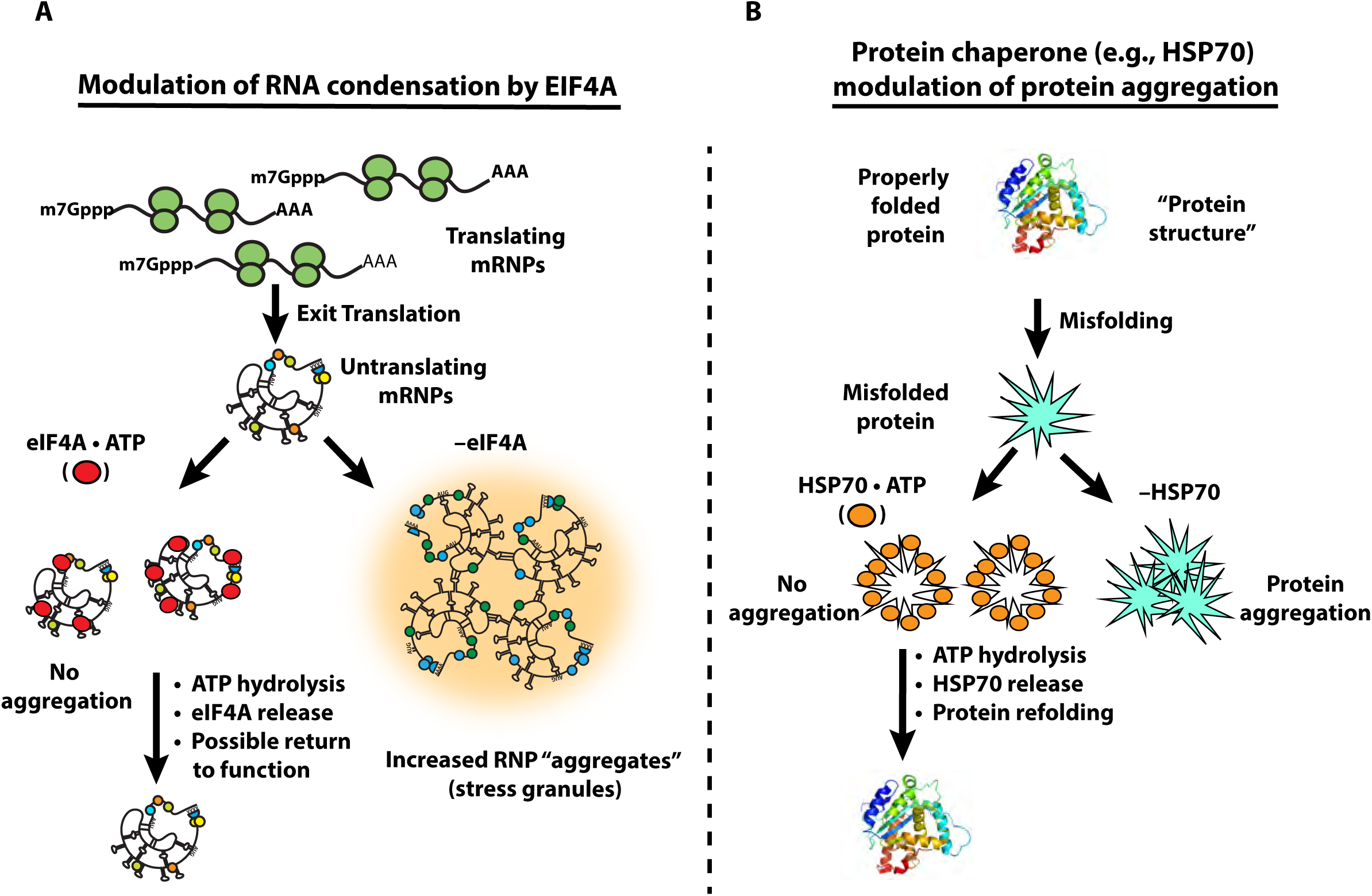
eIF4A limits RNA condensation as an ATP-dependent RNA chaperone, analogous to heat shock proteins. (A) Our mechanistic model of eIF4A’s function to resolve aberrant RNA-RNA interactions. ATP-dependent binding of eIF4A to free RNA limits multivalent RNA-RNA interactions driving RNA condensation while ATP hydrolysis facilitates eIF4A release to re-enter the catalytic cycle. (B) The general mechanism utilized by chaperones like HSP70 to resolve aberrant protein-protein interactions. Misfolded proteins are bound by HSP70•ATP, which limits aggregation while ATP hydrolysis liberates the protein.

Since overexpressing ATPase-defective DEAD-box mutants is sufficient to prevent SGs (Figure 4 and S5), it is likely that high concentrations of monovalent RNA binding proteins can also reduce the condensation of RNA. This is consistent with the observation that overexpression of the abundant RNA binding protein YB-1 can limit the formation of SGs and the accumulation of certain transcripts in them, consistent with its RNP chaperone abilities *in vitro* (Bounedjah et al., 2014; Tanaka et al., 2014). Cells appear to have adapted to make use of this mechanism across domains of life; for instance, the bacterial cold shock protein CspA is present at higher intracellular concentrations than eIF4A (∼30 µM) and acts to ameliorate global RNA structure during the cold shock response, possibly including intermolecular RNA-RNA interactions (Zhang et al., 2018).

A general role for eIF4A in limiting RNA condensation in the cytosol provides an answer to the decades-long question of why eIF4A is such an abundant protein and present at ∼10X higher concentrations than other translation initiation factors (Figure 3A; Pause et al., 1994). Since ATP depletion is more effective at rescuing SG formation in cell lines lacking G3BP (Figure 4A), and has a larger effect on mRNA recruitment to SGs during arsenite stress (Figure 3C), we suggest that additional ATP-dependent mechanisms, potentially including other DEAD-box proteins found associated with SGs (Figure 3A), also limit RNP condensation into SGs. Consistent with other DEAD-box proteins limiting SG formation, we demonstrate that over-expression of DDX19A, which is closely related to eIF4A (Cencic and Pelletier, 2016), can limit SG formation without preventing translational inhibition (Figure S4E-G). Interestingly, knockout of the DHX36 helicase has been shown to lead to increased SG formation, but this appears to be due to constitutive activation of PKR and increased translation repression (Sauer et al., 2019). An important area of future work will be identifying additional helicases that limit RNA condensation and determining if they act in RNA specific manners.

The generality of RNA condensation predict it will be relevant to other RNP granules. Indeed, observations in the literature are consistent with other RNA helicases functioning to limit RNAs being retained in RNA condensates. For example, knockdown of the RNA helicase UAP56/DDX39B (which is also related to eIF4A; Cencic and Pelletier, 2016), leads to increased accumulation of polyA^+^ RNA or fully spliced beta-globin mRNA in nuclear speckles (Dias et al., 2010), which are RNP condensates formed near sites of transcription. Similarly, catalytic UAP56 mutations lead to an accumulation of influenza M mRNAs in speckles (Hondele et al., 2019). Additionally, knockdown of the UPF1 RNA helicase, leads to retention of nascent transcripts in nuclear foci with DNA (Singh et al., 2019), where a high local concentration of nascent RNA might lead to RNA entanglements. These results suggest that RNA chaperones may be generally required to prevent the trapping of RNAs in thermodynamic energy wells at RNP granules. Additional mechanisms cells could utilize to limit RNA condensation include ribosome association, modulating RNA concentrations through RNA decay (Burke et al., 2019) or synthesis rates, and RNA modifications that alter the stability of RNA-RNA interactions.

A role for eIF4A, and other general RNA helicases, in limiting RNA condensation can be considered analogous to protein chaperones, such as HSP70, limiting the aggregation of misfolded proteins (Figure 7A&B). Multiple protein chaperones, including HSP70 proteins, bind to protein aggregates to disassemble aberrant interactions, thereby allowing for protein refolding and the solubilization of protein aggregates, using ATP hydrolysis as a switch for binding (Mogk et al., 2018). We suggest that RNA condensation and inappropriate aggregation occurs when the amount of exposed RNA in the cell exceeds the capacity of the cellular machinery limiting RNA condensation. Thus, the intrinsic aggregation properties of both proteins and RNAs are countered by abundant cellular machinery to keep these macromolecules correctly folded and dispersed for proper function.

## Acknowledgements

We thank the Parker lab, Amy Buck, and Olke Uhlenbeck for discussions. We thank the University of Colorado BioFrontiers Institute Cell Culture Facility and the BioFrontiers Institute Advanced Light Microscopy Core Facility (NIST-CU cooperative agreement 70NANB15H226 and NIH 1S10RR026680-01A1), John Rinn and Matt Disney for providing reagents. mCherry-DDX19A plasmids were a kind gift from Yaron Shav-Tal. This work was funded by NSF SCR training grant T32GM08759 (to D.T.), a Banting Postdoctoral Fellowship (to A.K.), NIH GM045443 (to R.P.) and the Howard Hughes Medical Institute.

## Author contributions

D.T., G.T., B.VT., A.K., and R.P. conceived the project. D.T., G.T., B.VT. and A.K. designed experiments, performed, and analyzed experiments. J.P. provided intellectual input and materials. D.T., G.T., and R.P. wrote the manuscript.

## Data and materials availability

All data is available in the manuscript or supplementary materials.

## Supplemental Figure Legends

**Figure S1. Recruitment of RNAs to RNA homopolymer condensates.** Related to Figure 1. (A) Pairwise combinations of fluorescent RNA homo-oligonucleotides and homopolymer RNA scaffolds were condensed together. Different oligo-scaffold combinations show differential recruitment due to differential RNA-RNA interaction strengths. Watson-Crick interactions promote strong, specific oligo recruitment, while noncanonical interactions can drive a range of partitioning strengths Some oligos display surface localization, suggesting that they homotypically interact while reducing surface free energy. Inset scale bars are 3 µm. (B) Quantification of (A) as indices of dispersion. Error bars represent 95% confidence intervals. *n* = 5 frames per condition. (C) Image displaying homopolymer droplets interacting following a physical disruption, indicating that heterotypic droplets physically interact. (D) Recruitment of fluorescent *in vitro* transcribed RNAs to homopolymer condensates. All RNAs tested were robustly recruited to the surfaces of polyA and polyU condensates. In contrast, some RNAs were recruited to the surfaces of polyC condensates, whereas others displayed punctate internalization, indicative of transcript RNA self-assembly within the polyC condensate.

**Figure S2. FRAP of polyA/*pgc* co-condensates.** Related to Figure 2. (A) Representative images of polyA/*pgc* co-condensates (with polyA visualized by fluorescent U_19_) before photobleaching and during recovery. (B) FRAP of polyA/*pgc* co-condensates. While the U_19_ signal recovers, *pgc* displays virtually no recovery, indicating that the surface *pgc* does not exchange. Fluorescent U_19_ recovery was fit to a one-phase association exponential curve (see STAR Methods; calculated *t*_1/2, oligoU_ ∼ 20 min), while *pgc* could not be fit due to the lack of recovery. (C) Representative images showing fluorescent recovery of polyA/*pgc* co-condensates following partial photobleaching. (D) Quantification of (C). *pgc* signal does not recover, indicating that *pgc* does not diffuse within the surface shell. Error bars represent ± 1 standard deviation of the mean. *n* ≥ 6 droplets. Error values for mobile fractions are calculated standard deviations.

**Figure S3. eIF4A inhibition increases partitioning of mRNA and G3BP into SGs.** Related to Figure 3. (A) Representative smFISH images displaying specific RNA localization (red) to SGs (green, G3BP) (B) Quantification of images in (A). (C) Representative G3BP1 IF and oligo(dT) FISH images in cells treated with arsenite, hippuristanol, or PatA. All images have brightness and contrast set to the same scale and are displayed as merged images and heatmaps. Scale bars are 5 µm. Hippuristanol treated cells demonstrate enhanced G3BP1 and polyA^+^ RNA partitioning to SGs. (D) Quantification of G3BP1 and polyA^+^ recruitment to SGs. Top: G3BP1 and polyA^+^ partition coefficients in cells treated with arsenite (A), hippuristanol (H), or PatA (P). Bottom: Fraction of G3BP1 or polyA^+^ in SGs in each condition. (E) Quantification of average SG area under arsenite, hippuristanol, or PatA, demonstrating that eIF4A inhibition increases the density of SGs, presumably by increasing RNA partitioning and RNA-RNA interactions. \**p*<0.05, \*\**p<*0.01, \*\*\**p*<10^−3^, *n* = 3 replicates.

**Figure S4. Effects of eIF4A inhibition, knockdown or over-expression on SGs are independent of translation, can be recapitulated with DDX19A, and do not alter RNA binding protein exchange dynamics.** Related to Figure 4. (A) Immunoblot depicting puromycin labeling of ΔΔG3BP1/2 cells treated with arsenite (500 µM), hippuristanol (1 µM), PatA (100 nM), or combinations of arsenite and hippuristanol or PatA. In all conditions, translation is inhibited >97%, indicating that additional translational inhibition with combinatorial drug treatments is not responsible for SG recapitulation. (B) Immunoblot depicting siRNA mediated knock-down of eIF4A1 showing decreased translation as expected for depletion of a translation initiation factor. (C) Immunoblot depicting Myc-eIF4A1 OE. Note that OE of eIF4A1 does not increase bulk levels of translation, indicating that prevention of SG formation upon arsenite addition is not due to increased basal translation. (D) Puromycin IF displaying translational inhibition in cells containing over-expressed Myc-eIF4A1 after arsenite addition, indicating that eIF4A1 OE does not prevent translational shutoff. (E) Overexpression of mCherry-DDX19A can limit arsenite-induced SG formation (quantified to the right by SG area as a percentage of cell area [%SG area/cell area]), and these effects were not observed with mCherry expression alone, consistent with other helicases containing the eIF4A helicase core sequence functioning as RNA chaperones. ATPase defective mutant (E242Q) was able to repress SG formation similar to Myc-eIF4A1 E183Q, while RNA binding mutant E372G was not. This data is consistent with RNA binding being required for DEAD-box helicases to function as RNA chaperones (*see also* Figure 4). (F) Puromycin IF depicting translational shutoff in cells over-expressing mCherry-DDX19A, confirming that DDX19A does not prevent translational shutoff caused by arsenite. (G) Quantifications of SG formation from mCherry-DDX19A over expression experiments depicted in (E). (H) Quantification of GFP-G3BP1 FRAP between arsenite (500 µM), PatA (100 nM), or hippuristanol (1 or 4 µM) showing no differences in the exchange between stressors, indicating that eIF4A inhibition does not influence the exchange rate of G3BP1 in SGs. For all quantifications, error bars represent standard deviations, *n* ≥ 3 independent replicates.

**Figure S5. Δ/ΔG3BP1/2 SGs contain similar RNA and protein compositions and properties to wildtype.** Related to Figure 4. (A) SGs that form in Δ/ΔG3BP1/2 cells due to arsenite combined with eIF4A inhibition contain canonical SG proteins: PABPC1, TIA1, eIF4G, eIF4A1, and eIF4B. SGs do not overlap with PB marker EDC3, similar to wildtype cells, and SG formation is inhibited with cycloheximide (CHX) pre-treatment, indicating that ΔΔG3BP1/2 SGs require non-translating mRNAs to form. (B) Hippuristanol and arsenite Δ/ΔG3BP1/2 SGs contain the lncRNA *AHNAK*, (C) polyadenylated RNA and the SG-enriched lncRNA *NORAD*.

**Figure S6. Representative images and quantifications of RNA condensation inhibition control experiments.** Related to Figure 6. (A) Images showing the effects of RNA alone, ATP (1 mM), ADPNP (1 mM), ATP + 10 µM hippuristanol, or ATP + 1 µM pateamine A on total RNA droplet formation. (B) quantifications of (A). No significant changes in droplet formation were observed between ATP, ADPNP, or drug combinations. *n* = 3 for each experiment and error bars represent standard deviations of the mean. SB = Protein storage buffer.

**Movie S1. Time-lapse microscopy reveals the growth of polyC/polyU RNA condensate networks.** Related to Figure 1. Maximum intensity projections of deconvoluted 2 µm Z-stacks of polyC (green) and polyU (red) condensates, labeled by fluorescent antisense RNA oligonucleotides, over time. Each frame represents 3 minutes of real time, with the first frame occurring 3 minutes following mixing with condensation buffer. Note that heterotypic droplets are associated at early timepoints, and that this association scaffolds the growth of the network.

**Movie S2. Time-lapse spinning disk confocal microscopy of polyA/*pgc* co-condensates following dilution.** Related to Figure 2. PolyA, labeled with fluorescent U_19_ (magenta), was condensed with fluorescent *pgc* (green) and thereafter subjected to 1:10 dilution in TE buffer, dissolving the polyA assemblies while the *pgc* assemblies persist for many minutes. Scale bar is 20 µm. Each frame represents 3 seconds of actual time.

## STAR Methods

### KEY RESOURCES TABLE

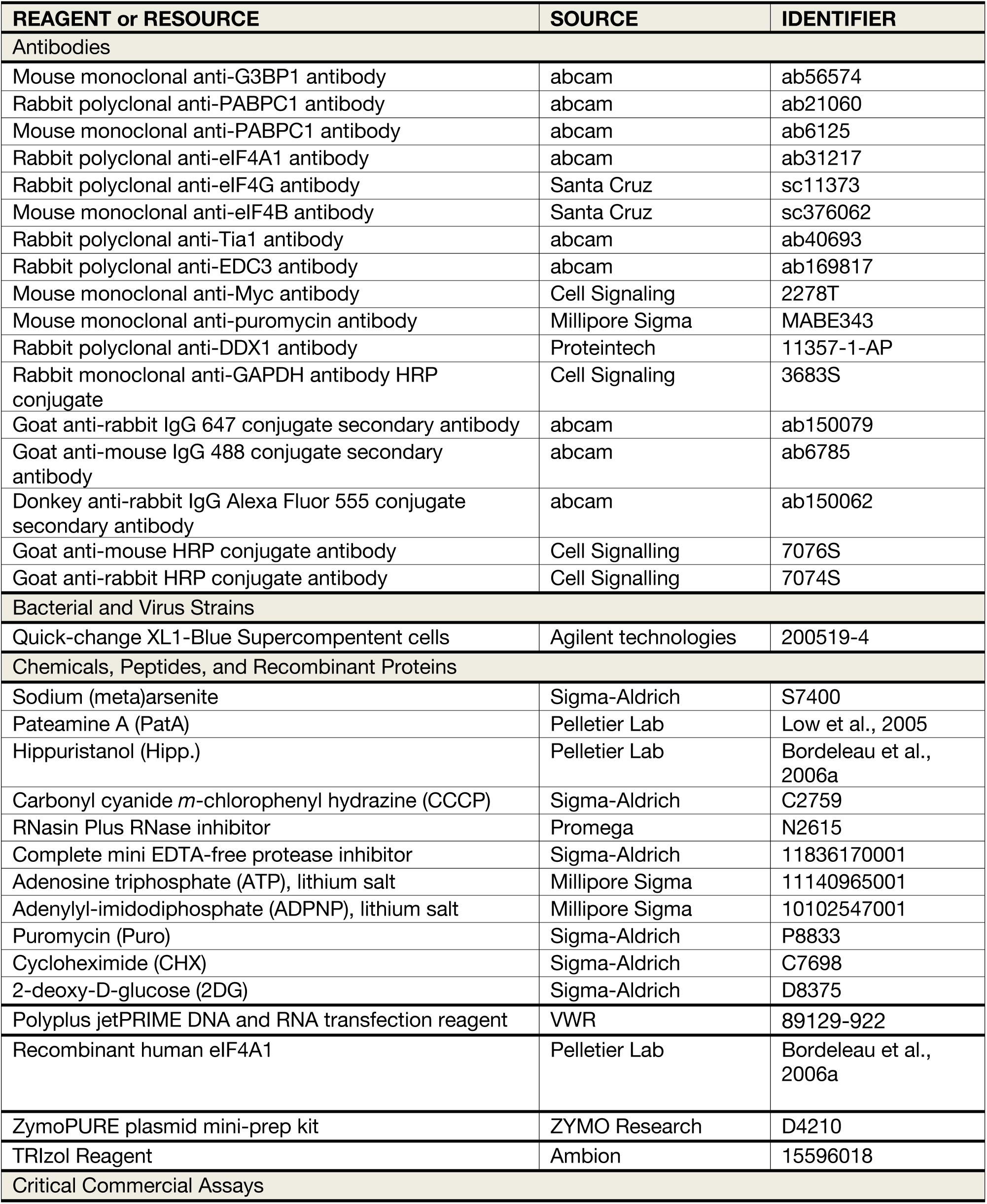
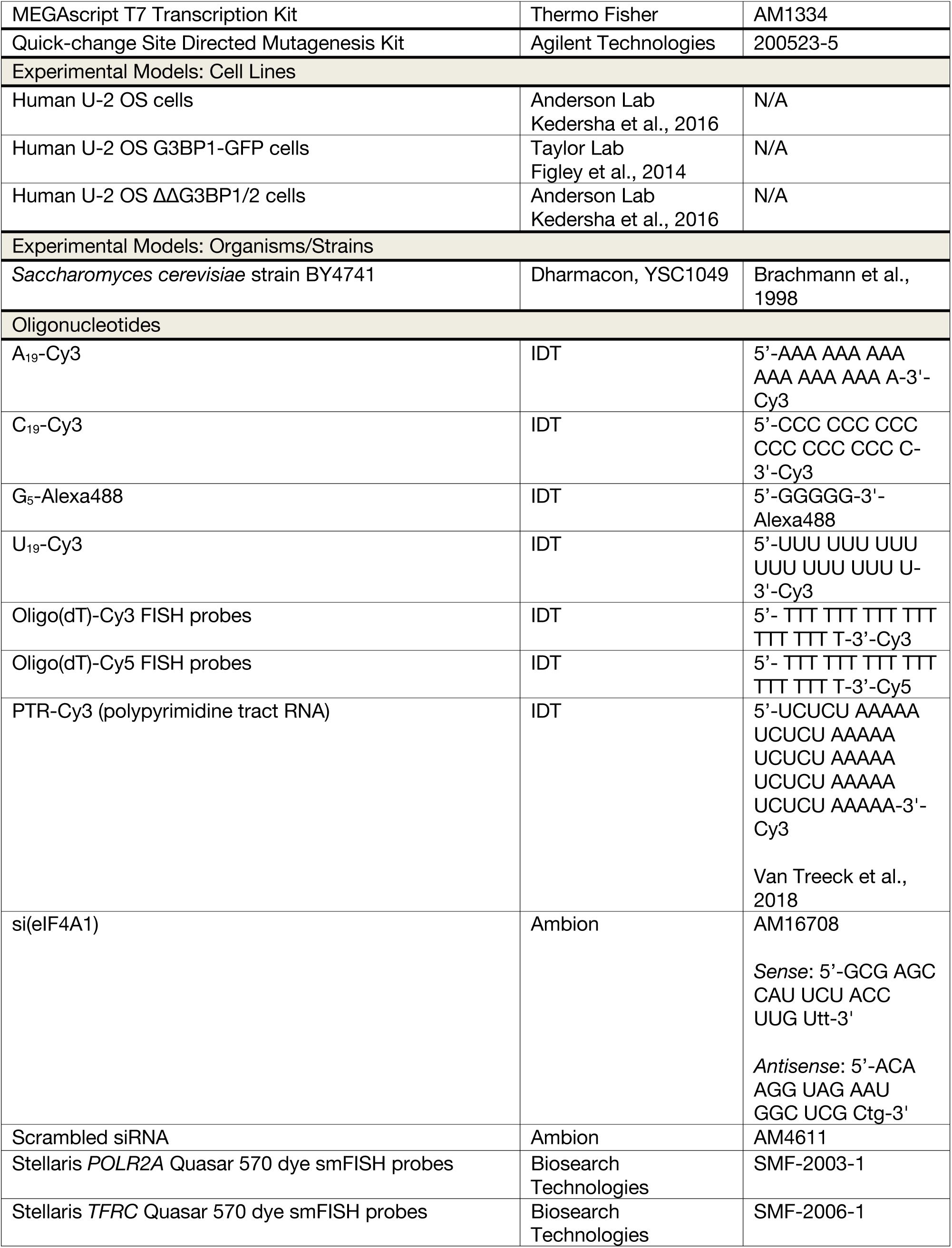
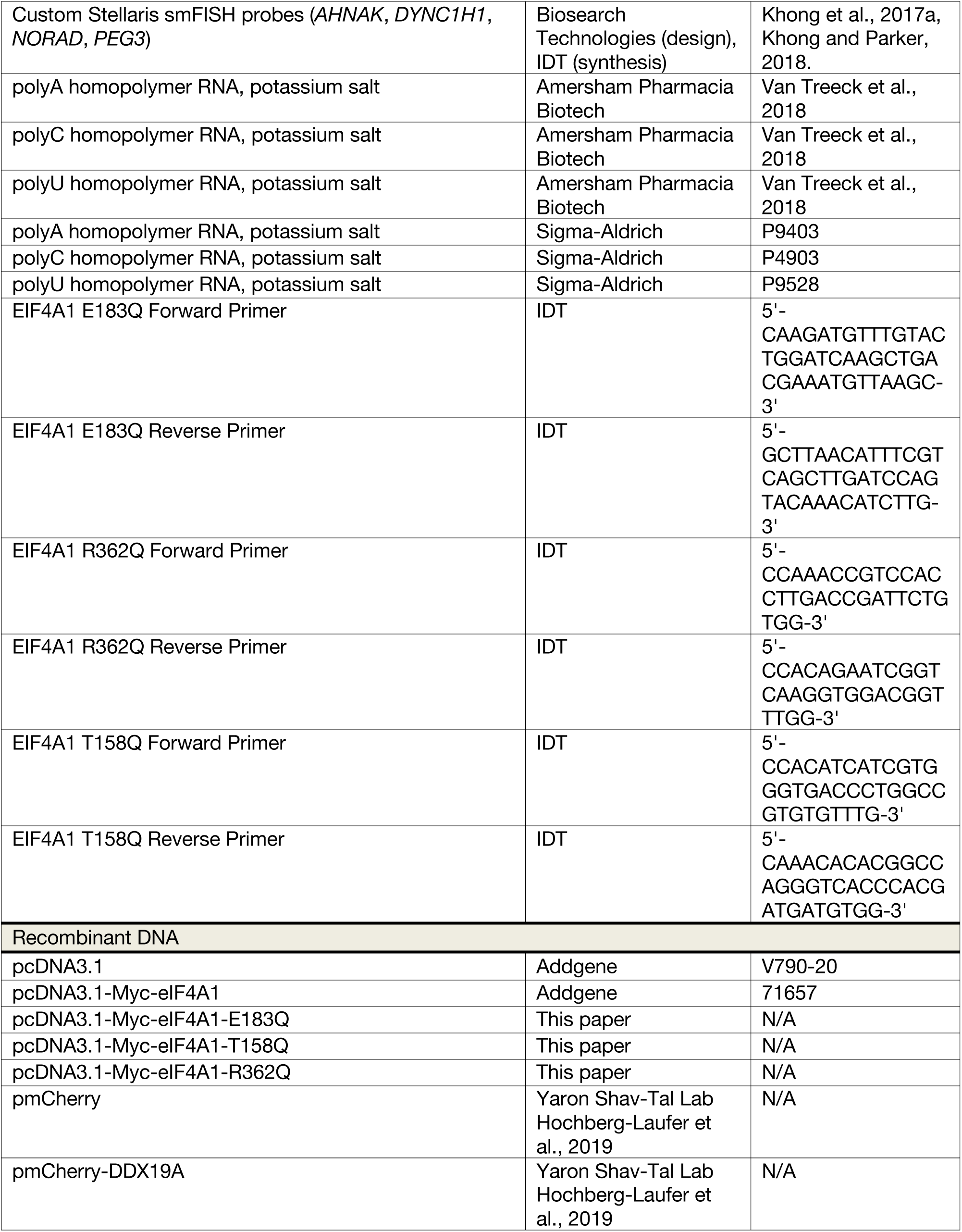
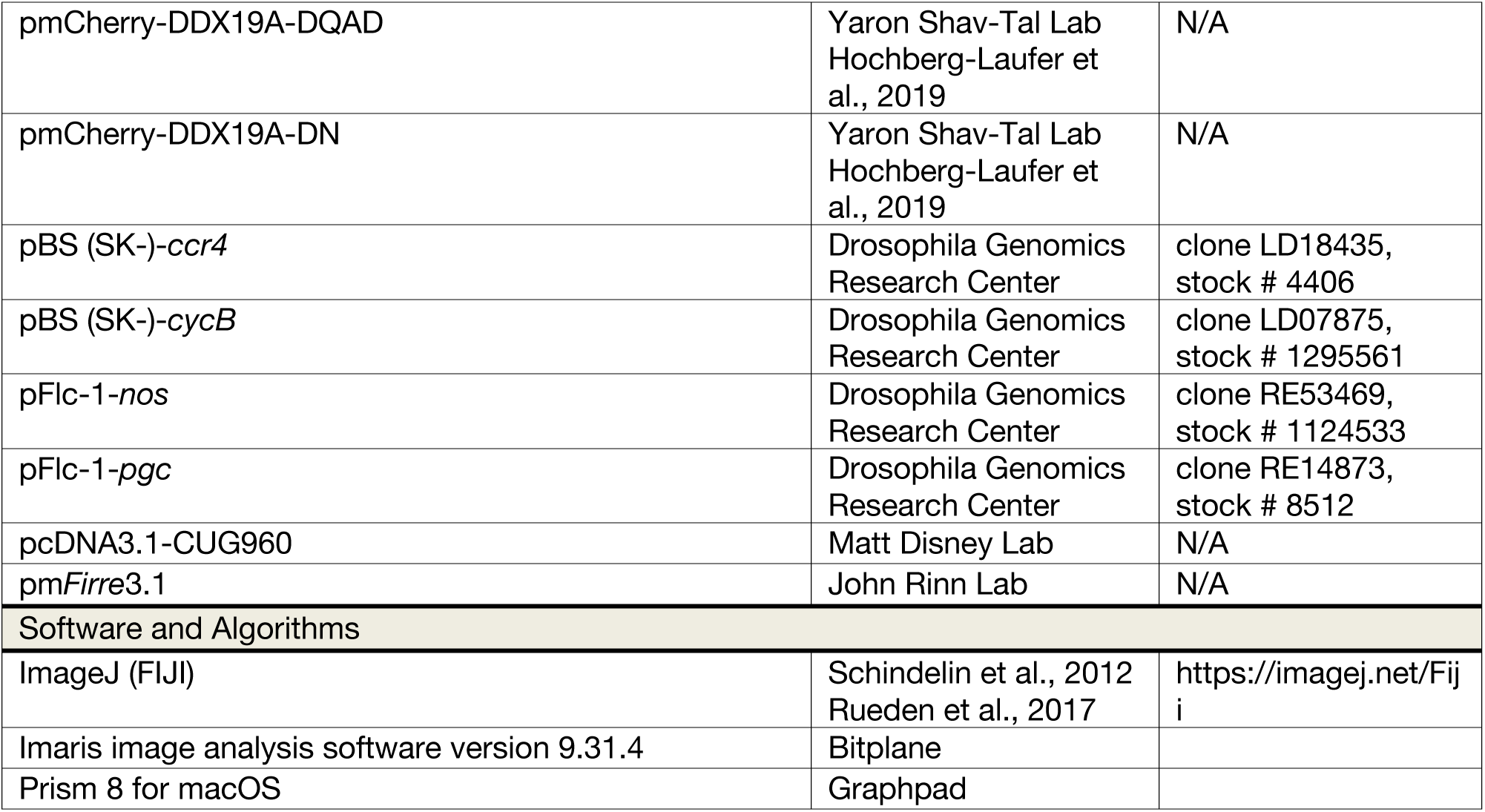

### CONTACT FOR REAGENT AND RESOURCE SHARING

Further information and requests for resources and reagents should be directed to and will be fulfilled by the lead contact, Roy Parker.

### EXPERIMENTAL MODEL AND SUBJECT DETAILS

#### Cell lines and growth conditions

Cell lines used in this study include WT parental human U-2 OS cells, ΔΔG3BP1/2 double knockout U-2 OS cells, and G3BP1-GFP U-2 OS cells. U-2 OS cells are female. All cell lines were grown and maintained at 37°C and 5% CO_2_. Cells were grown in Dulbecco’s modified Eagle’s medium (DMEM) with 10% fetal bovine serum (FBS) and 1% penicillin/streptomycin mix.

#### Yeast and bacterial strains and growth conditions

For total RNA extraction, *Saccharomyces cerevisiae* strain BY4741 (Dharmacon) was cultured in yeast extract peptone dextrose (YEPD) media at 30°C. For plasmid isolation, XLI-Blue supercompetent cells were grown in lysogeny broth (LB) at 30°C.

### METHOD DETAILS

#### Plasmid Isolation and Site-Directed Mutagenesis

Plasmids were amplified by transformation into XL1-Blue supercompetent cells (Agilent Technologies) by following manufacturer’s instructions. Bacteria was then grown overnight in 5 mL cultures in LB media with respective selectable marker antibiotic. Plasmids were then isolated using ZymoPURE plasmid mini-prep kit by following manufacturer’s instructions. Plasmid concentrations were determined via absorbance at 260 nm using a Nanodrop 2000 device (Thermo Scientific). Site directed mutagenesis was performed on pcDNA3.1-Myc-eIF4A1 by using the Quick-change Site Directed mutagenesis kit (Agilent Technologies) according to manufacturer’s instructions. Mutations were verified via commercial sanger sequencing (Genewiz).

#### *In vitro* transcription of RNAs

*Drosophila nos*, *pgc*, *ccr4*, and *cycB* T7 transcription plasmids were procured from the Drosophila Genomics Research Center (DGRC; Bloomington, IN). The m*Firre*3.1 T7 transcription plasmid was generously provided by the John Rinn Lab at CU Boulder. The CUG repeat RNA construct was generously provided by the Matt Disney Lab (Scripps Research Institute). Plasmids were linearized via restriction digestion with BamHI-HF for the *Drosophila* RNAs, Afe1 for luciferase, and Kpn1-HF for *Firre* (enzymes from New England Biolabs (NEB)). The CUG repeat construct was linearized with HindIII-HF (NEB). Restriction digestion reactions were performed according to the manufacturers’ recommendations. Following linearization, plasmid DNA was recovered with ethanol precipitation followed by resuspension in TE buffer (10 mM Tris HCl pH 8.0, 1 mM EDTA).

*In vitro* transcription of fluorescent RNAs was performed using the T7 MEGAscript kit (Ambion) according to the manufacturer’s recommendations with the following modification: Addition of unlabeled UTP was reduced by 25% and an equal amount of fluorescently labeled UTP (fluorescein-12-UTP or cyanine-5-UTP, Enzo Life Sciences) was added into the reaction mixture. Following transcription, DNA was removed by treating with TURBO DNase I (Ambion) according to the manufacturer’s recommendations. RNA was recovered by sequential acid phenol/chloroform and chloroform extractions, followed by precipitation in isopropanol and NH4OAc. RNA recovery was quantified via UV-Vis at 260 nm using a Nanodrop 2000 device (Thermo-Fischer). Proper RNA sizes were validated through either denaturing gel electrophoresis or Agilent TapeStation RNA ScreenTape analysis (performed by the BioFrontiers Next-Generation Sequencing Core Facility). Recovered RNA was resuspended in RNase-free water (Invitrogen) or TE and stored at –80°C. Working 100 μg/ml stocks were created as needed by diluting appropriately into RNase-free water and kept at –80°C.

#### Homopolymer condensation

Homopolymer RNAs were purchased as salts from Amersham Pharmacia Biotech, Inc. (polyA, polyC, and polyU) and Sigma-Aldrich (polyU) and made into 25 mg/mL stock solutions in TE or RNase-free water, then kept at –80°C. Working 5 mg/mL stock solutions were prepared by diluting into RNase-free water. These working stocks were stored at –80°C or –20°C. The fluorescently-labeled oligonucleotides were purchased from IDT and resuspended, from which 2 μM stock solutions were made and stored at –80°C. An initial aliquot of the fluorescently-labeled PTR (*p*olypyrimidine *t*ract *R*NA; Table S1) was purchased from IDT and was stored at –80°C as a 2 μM stock solution.

To form homopolymer condensates for the short RNA localization experiments, homopolymer RNA was condensed as described previously (Van Treeck et al., 2018), with the addition of 200 nM fluorescent RNA oligos in the condensation mix. To localize *in vitro* transcribed RNAs, homopolymer RNA was condensed with the transcribed RNA in the same manner as above, but instead mixed with 150 mM NaOAc, 600 mM NaCl, 1 mM MgCl_2_, and 10% w/v PEG 3350 (all in RNase-free water) to a final homopolymer concentration of 0.4 mg/ml and a final mRNA/lncRNA concentration of 10 μg/ml, yielding an approximate pH of 7 as assessed by pH paper.

#### Mechanical disruption of polyC/polyU networks

To mechanically disrupt polyC/polyU condensate networks, condensates were first prepared in glass bottom 96-well plates and incubated for an hour. Thereafter, condensates were pipetted into adjacent empty wells, and pipetted up and down quickly five times. Afterwards, they were immediately imaged.

#### Repeat RNA (reRNA) condensation

CUG repeat foci were prepared by condensing 200 μg/ml of fluorescein-labeled *DMPK*^Exons 11-15^-CUG_590_ RNA as in Jain and Vale (2017), in the presence or absence of 10 μg/ml Cy5-labeled *in vitro* transcribed RNAs or 200 nM PTR-Cy3. Briefly, RNAs were mixed together in a buffer of 10 mM MgCl_2_, 10 mM Tris HCl pH 6.8, and 25 mM NaCl, and subsequently denatured for 3 min at 95°C then cooled at ∼4°C/min to 37°C in a thermocycler. Afterwards, condensates were immediately imaged.

#### Stress granule isolation and RNA recruitment

To test the ability of SGs to recruit *in vitro* transcribed RNAs, SGs were purified as follows: First, GFP-G3BP1 U-2 OS cells were seeded at ∼40% confluency and then grown to ∼80% confluency, after which the media was replaced and the cells were stressed with 500 μM arsenite for 60 min. Afterwards, the cells were washed, pelleted, and flash-frozen in liquid N2, then stored at –80°C for up to a week. SGs were isolated as in Khong et al. (2017b) to produce SG-enriched fraction. No affinity purification was performed. Other cellular debris was removed from the SG-enriched supernatant through both vigorous pipetting and by passing the supernatant through a 25G 5/8 needle, accompanied by quick spins. RNA recruitment to SGs was assessed by incubating 10 µg/mL Cy5-labeled luciferase RNA with SGs for 10 minutes at room temperature, followed by imaging.

#### RNA condensate dilution assays

To test the stability of RNA assemblies under dilution, *pgc*-decorated polyA droplets were prepared as above and incubated 2 hours. Thereafter, they were subjected to spinning disk confocal time-lapse microscopy. Every 18 sec, an image was taken in each channel in a 4×4 grid of adjacent frames using the Nikon Elements software “Large Image” tool. These images were automatically stitched together. Dilutions were performed while imaging was ongoing. In between imaging timepoints, TE buffer was quickly added dropwise (so as to only minimally disturb assemblies) to a tenfold final dilution. The very next imaging timepoint was taken to be *t* = 0 for quantitative analysis.

#### RNA crosslinking assays

To observe RNA-condensate-associated RNA-RNA interactions, 4’-aminomethyltrioxsalen (AMT) UV crosslinking was performed (Frederikson and Hearst, 1979). AMT was procured from Santa Cruz Biotechnology and solubilized to a final concentration of 1 mg/mL in DMSO. AMT stocks were stored at –20°C. Since AMT crosslinks pyrimidine residues, condensation-crosslinking experiments were performed using polyA condensates to avoid crosslinking the client RNA to the homopolymer scaffold.

In condensation-crosslinking experiments, RNA was demixed in the appropriate condition, but with the addition of 100 μg/ml AMT. Thus, droplets and tanglets were crosslinked in 600 mM NaCl, 150 mM NaOAc, 1 mM MgCl_2_, 10% PEG MW 3350, 0.1 mg/mL AMT. Crosslinking in solution conditions was performed in TE with 0.1 mg/mL AMT added. Following incubation, RNA was crosslinked by irradiating 15 min with 366 nm UV light. Afterwards, samples were diluted 1:1.7 in TE and RNA was recovered by precipitation in 2 M LiCl. If RNA sample concentrations were low (e.g., tanglet conditions), 0.4 mg/ml polyA homopolymer RNA was added post-crosslinking and pre-precipitation as a carrier in order to promote efficient recovery.

#### Denaturing gel electrophoresis

Denaturing gel electrophoresis was performed to analyze fluorescent and crosslinked RNAs. RNA samples were mixed with either 6x formaldehyde loading dye (Ambion) or 2x formamide loading dye (95% deionized formamide, 0.025% w/v bromophenol blue, 5 mM EDTA) and denatured by incubation at 70°C for 15 min, followed by incubation on ice for 3 min. Samples were run in freshly-prepared 1% agarose, 1% formaldehyde denaturing gels in fresh 1x MOPS buffer (20 mM MOPS pH 7.0, 5 mM NaOAc, 1 mM EDTA) at 4-10 V/cm for 2.5-4 hours.

Denaturing gels were imaged using a Typhoon FLA 9500 imaging system (GE) set to visualize the appropriate fluorescence signal from fluorescently-labeled RNAs. Crosslinking efficacies were quantified in ImageJ by measuring the integrated intensities of the monomer bands and higher-weight bands, subtracting the average of 10-15 background measurements, and dividing the sum of the corrected higher-weight integrated intensities by the sum of all band integrated intensities. In symbols:

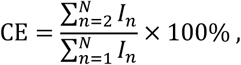

where *I_n_* is the (background-corrected) integrated intensity of the band corresponding to the RNA *n*-mer, *N* is the highest order band observed in a given sample, and CE is the crosslinking efficacy.

#### Total RNA isolation and *in vitro* decondensase assays

*Saccharomyces cerevisiae* strain BY4741 was grown to an OD600 of 0.8 AU and was lysed with 425-600 µm glass beads (Sigma-Aldrich) and vortexing for 10 minutes in a 50 mL conical tube in TRIzol (Ambion). RNA was extracted by using TRIzol reagent according to manufacturer’s instructions. Purified RNA concentration was determined via absorbance at 260 nm using a Nanodrop 2000 device (Thermo Scientific).

To assess the ability of eIF4A1 to decondense RNA *in vitro*, 10 µM recombinant eIF4A1 or protein storage buffer (200 mM NaCl, 25 mM Tris, 10% glycerol, 1 mM DTT, pH 7.4) was mixed together in with 150 µg/mL yeast total RNA extracts, SYTO^TM^ RNAselect^TM^ green fluorescent dye, and either 1 mM ATP or ADPNP (Sigma-Aldrich) in 150 mM KOAc, 1 mM MgCl_2_, 0.223 mM spermine tetrahydrochloride, 1.34 mM spermidine trihydrochloride, and 8% w/v PEG 3350 in a glass bottom-96 well plate at 37°C. These conditions yield an approximate pH of ∼7.4. Polyamine levels recapitulate physiological concentrations (Van Treeck et al., 2018). Drug treatments were performed with DMSO or 10 µM hippuristanol or 1 µM pateamine A. For each reaction, RNA condensate formation was observed after initial RNA addition by DIC and FITC channels over 30 minutes with images taken every 5 minutes. %Droplet area was analyzed using “Particle finder” in ImageJ.

#### Cell drug treatments and transfections

For drug treatments or transfections, cells were seeded at ∼10^5^ cells/mL and were allowed to adhere overnight. For siRNA experiments, cells were transfected with 50 nM of scrambled or eIF4A1 siRNA (see Table S1) using Interferin (polyPLUS) according to the manufacturer’s instructions for 48 h before SG induction. For overexpression experiments, cells were transfected with control (pcDNA3.1, Addgene) or pcDNA3.1-Myc-eIF4A1 (Addgene) plasmids for 48 h prior to SG induction or pmCherry (control) or pmCherry-DDX19A plasmids for DDX19A overexpression.

To induce SGs, arsenite (500 μM in H_2_O, Sigma-Aldrich), hippuristanol (Hipp., 300 nM or 1 µM in DMSO), or Pateamine A (PatA, 100 nM in DMSO) were added and cells were incubated at 37°C for the allotted times indicated in each assay. To deplete ATP, cells were incubated with 2-deoxy-D-glucose (2DG, 200 mM in H_2_O, Sigma-Aldrich) and carbonyl cyanide *m*-chlorophenyl hydrazine (CCCP, 100 μM in DMSO, Sigma-Aldrich). For ribopuromycinylation assays, cells were incubated at 37°C with puromycin (10 μg/mL in H_2_O, Sigma-Aldrich) 5 minutes prior to fixation or lysis.

#### Immunoblotting

Following drug treatment or transfection, cells were washed with 37°C phosphate buffered saline (PBS) and lysed with NP-40 lysis buffer (50 mM Tris-HCl pH 8.0, 150 mM NaCl, 1% NP-40 and protease inhibitor cocktail (Thermo Scientific)). Cell lysates were rocked at 4°C for 30 minutes, and then clarified by centrifugation (13K RPM for 60 s). 4x Nu-PAGE sample buffer was added to lysates to a final concentration of 1x, samples were boiled for 5 minutes at 95°C, and then loaded into 4-12% Bis-Tris Nu-PAGE gel and transferred to a nitrocellulose membrane. Membranes were blocked with 5% BSA in Tris-buffered saline with 0.1% Tween-20 (TBST) for an hour and then incubated with primary antibody overnight at 4°C. Membranes were washed 3x with TBST, then incubated at room temperature for 2 hours in 5% BSA in TBST. Membranes were washed 3x again in TBST and antibody detection was achieved by rocking membranes in Pierce ECL western blotting substrate for 5 minutes.

Chemiluminescence was visualized on an Image Quant LAS 4000 (GE). Protein band densities were quantified in ImageJ.

#### Immunofluorescence (IF) and fluorescence *in situ* hybridization (FISH)

Cells were prepared as described as stated above except grown on No. 1.5 glass coverslips (Thermo). Sequential IF/FISH or IF/smFISH was performed as previously described (Khong et al., 2017a; Khong et al., 2017b). When IF was performed without FISH, fixed U-2 OS cells were instead blocked with 5% BSA in PBS for 1 hour prior to IF and the antibodies were diluted in 5% BSA in PBS.

Alternatively, fixed cells were simultaneously blocked and permeabilized with 5% BSA in PBS-T (0.1% Triton-X-100) for 1 hour at room temperature. Thereafter, coverslips were incubated with primary antibody (1:500) overnight at 4°C in 1% BSA in PBS-T. Coverslips were then washed 3X with PBS and incubated with secondary antibody (1:1000) at room temperature for 2 hours in 1% BSA in PBS-T.

The primary antibodies used for immunofluorescence (IF) include mouse anti-G3BP1 (5 µg/mL, ab56574(Abcam)), rabbit anti-DDX1 (1:100, 11357-1-AP (Proteintech)), and rabbit anti-eIF4A1 (100 µg/mL, ab31217(Abcam)) and the appropriate secondary antibodies used were goat anti-mouse FITC antibody (1:1000, Abcam (ab6785)), and donkey anti-rabbit Alexa Fluor 555 conjugate antibody (1:500, Abcam (ab150062).

*NORAD*, *PEG3*, and *DYNH1C1* smFISH probes were custom made using Stellaris® RNA FISH probe Designer (Biosearch Technologies, Inc., Petaluma, CA). The probe sequences are listed in Table S1. *AHNAK*-Quasar670, *TFRC*-Quasar570, and *POLR2A*-Quasar570 probes were purchased premade directly from Biosearch Technologies and resuspended according to the manufacturer’s recommendations. *AHNAK* smFISH probe sequences are in Khong et al. (2017a). Oligo(dT)-Cy3 probes were purchased from IDT.

#### Microscopy

Fixed U-2 OS cells stained by IF and/or smFISH, purified SGs, CUG repeat RNA foci, and homopolymer condensates were imaged using a widefield DeltaVision Elite microscope with a 100x NA 1.5 oil objective using a PCO Edge sCMOS camera and SoftWoRx software (GE).

Condensate dilution and helicase assays were imaged using an inverted Nikon Ti Eclipse spinning disk confocal microscope equipped with an environmental chamber and Nikon elements software. Helicase assay imaging was performed at 37°C. All spinning disk confocal images were acquired with a 100x NA 1.5 oil objective and a 2x Andor Ultra 888 EMCCD camera.

#### Fluorescence recovery after photobleaching (FRAP)

FRAP assays were performed using an inverted Nikon A1R laser scanning confocal microscope equipped with an environmental chamber, a 100x NA 1.5 oil objective, and Nikon Elements software. At least three images were acquired prior to photobleaching followed imaging over the course of recovery. PolyA/*pgc* FRAP was performed at room temperature.

To analyze recovery, the mean intensity of each bleached region was quantified in ImageJ, and recovery intensities were normalized to the mean of three pre-bleach measurements. Mobile fractions *φ_M_* were computed by subtracting the minimum normalized mean intensity *I*_0_ from the normalized endpoint intensity *I_F_*: *φ_M_* = *I_F_* – *I*_0_. OligoU recovery in polyA droplets was modeled by a single association exponential curve using Prism Graphpad software.

Partial FRAP was performed similarly, but by only photobleaching approximately half the condensate area.

### QUANTIFICATION AND STATISTICAL ANALYSIS

Precision, dispersion, and *n* values are found and explained in the figure legends.

#### Image analysis

Single molecule FISH analysis was performed using Imaris (Bitplane) as in (Khong et al., 2017a; Khong et al., 2017b; Khong and Parker, 2018). Briefly, processed Z-stacks are opened and the nuclei masked using the DAPI channel. Thereafter, smFISH spots are identified in 3D using the “Spots” tool and the automatic parameters. SGs are similarly identified in 3D with the “Cells” tool, with 3D masks rendered. The number of smFISH spots in SGs is quantified as the number of spots in the 3D rendered SGs. Thus, the fraction of RNA molecules in SGs, the data presented, is this value divided by the total number of smFISH spots identified.

G3BP1, eIF4A1, and oligo(dT) partitioning were quantified using Imaris Imaging Analysis software (Bitplane). In order to measure partitioning, first, the nuclei were masked using the DAPI source channel. Second, individual cells were segmented by creating a region of interest (manually determined) around a single cell using the surface creation wizard. Mean cellular fluorescence intensities for G3BP1, eIF4A1, and oligo(dT) were then obtained under the results tab. Third, SGs were identified using the G3BP1 fluorescent source channel with Imaris Cell Creation wizard with the following parameters (0.0406 µm filter width, manually-determined thresholding, and *≥* 1 voxel). Once SGs were identified by the cell creation wizard, they are then converted to surfaces in order to extract mean fluorescence intensities for G3BP1, eIF4A1 and oligo(dT) inside stress granules. Partitioning was then calculated by taking the ratio of the mean fluorescence intensity in SG to the mean fluorescence intensity in the cell. 20 cells were counted for each sample. For more detailed information, please see Khong et al. (2017b).

Line analysis was performed using ImageJ (FIJI). A straight line was drawn across a stress granule using G3BP1 as a source channel. The line is then saved as an ROI. The ROI was then pasted on the corresponding eIF4A1 or DDX1 source channel. Line intensities were then extracted in both channels using the “Plot Profile” tool. Stress granule boundaries (in each source channel) were determined by an intensity threshold halfway between the maximum and minimum intensity values for each source channel. Fifty linescan analyses were performed for each experiment.

Measurements of SG area or PB area per cell area were quantified using the “Particle Finder” tool in ImageJ. The percent of cells with SGs was quantified using the “Count” tool in ImageJ. The fluorescence intensity of puromycin-labeled nascent peptides in individual cells was quantified as mean gray values extracted from ImageJ.

SG and PB interface calculations were quantified by identifying regions of overlap between each mRNP granule and recording these events by using the “count” tool in image J. Once total interfaces were counted, these values were normalized either to PB counts/area (determined using “analyze particles” in Image J) or SG counts/area (using “analyze particles” in ImageJ).

The index of dispersion (DI) is a statistical measure of inhomogeneity in a population and is defined to be

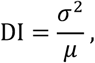

where σ^2^ is the variance of the population distribution (with σ being the population standard deviation) and *µ* is the mean value. For example, particles in an ideal solution exhibit a dispersion index of 1, as they are Poisson distributed, and substantially higher values indicate particle-particle interactions in that context. Here, dispersion indices were used to quantify inhomogeneity in fluorescence signal and infer the relative extent to which an RNA is assembled, similarly to Jain and Vale (2017).

Indices of dispersion were computed by measuring the mean fluorescence intensity and standard deviation for maximum projections of 1 µm Z-stacks or single slices in the case of the dilution experiment. An index was computed for each image using ImageJ, and the displayed index was taken to be the average of the indices for all the separate images. The standard deviations for the distribution of index values for a given set of conditions were used to calculate 95% confidence intervals.

#### Estimates of eIF4A1 levels per mRNA

For HeLa and U-2 OS cells we used published values for eIF4A1 from quantitative mass spectroscopy, 1.7×10^7^ and 2.2×10^6^, respectively (Itzhak et al., 2016; Beck et al., 2011), and mRNA levels for U-2 OS cells, ∼330,000 (Khong et al., 2017). Although previous estimates suggest HeLa cells might contain 200,000 mRNAs per cell (Shapiro et al., 2013) we utilized an estimate of 330,000 mRNAs per cell to be conservative in our estimations. These calculations indicate a ratio of 6.7 eIF4A1 molecules/mRNA to 51 eIF4A molecules/mRNA in U-2 OS and HeLa cells respectively.

#### Statistical analysis

Statistical significance was assessed using the Student’s *t*-test with two tails.

### DATA AND CODE AVAILABILITY

All data produced by this study are included in the manuscript or available from the authors upon request.

## References

Al-Husini, N., Tomares, D.T., Bitar, O., Childers, W.S., Schrader, J.M. (2018). α-Proteobacterial RNA Degradosomes Assemble Liquid-Liquid Phase-Separated RNP Bodies. Mol. Cell 71, 1027– 1039.

Anderson, P., and Kedersha, N. (2006). RNA granules. J. Cell Biol. 172, 803–808.

Aulas, A., Fay, M.M., Lyons, S.M., Achorn, C.A., Kedersha, N., Anderson, P., and Ivanov, P. (2017). Stress-specific differences in assembly and composition of stress granules and related foci. J. Cell Sci. 130, 927–937.

Aumiller, W.M., Jr., Pir Cakmak, F., Davis, B.W., and Keating, C.D. (2016). RNA-Based Coacervates as a Model for Membraneless Organelles: Formation, Properties, and Interfacial Liposome Assembly. Langmuir 32, 10042–10053.

Banani, S.F., Lee, H.O., Hyman, A.A., and Rosen, M.K. (2017). Biomolecular condensates: organizers of cellular biochemistry. Nat. Rev. Mol. Cell Biol. 18, 285–298.

Beck, M., Schmidt, A., Malmstroem, J., Classen, M., Ori, A., Szymborska, A., Herzog, F., Rinner, O., Ellenberg, J., and Aebersold, R. (2011). The quantitative proteome of a human cell line. Mol. Syst. Biol. 7, 549.

Bordeleau, M.E., Cencic, R., Lindqvist, L., Oberer, M., Northcote, P., Wagner, G., and Pelletier, J. (2006b). RNA-mediated sequestration of the RNA helicase eIF4A by pateamine A inhibits translation initiation. Chem. Biol. 13, 1287–1295.

Bordeleau, M.E., Mori, A., Oberer, M., Lindqvist, L., Chard, L.S., Higa, T., Belsham, G.J., Tanaka, J., and Pelletier, J. (2006a). Functional characterization of IRESes by an inhibitor of the RNA helicase eIF4A. Nat. Chem. Biol. 2, 213–220.

Bordeleau, M.E., Matthews, J., Wojnar, J.M., Lindqvist, L., Novac, O., Jankowsky, E., Sonenburg, N., Northcote, P., Teesdale-Spittle, P., and Pelletier, J. (2005). Stimulation of mammalian translation initiation factor eIF4A activity by a small molecule inhibitor of eukaryotic translation. Proc. Nat. Acad. Sci. USA 102, 10460–10465.

Bounedjah, O., Desforges, B., Wu, T.D., Pioche-Durieu, C., Marco, S., Hamon, L., Curmi, P.A., Guerquin-Kern, J.L., Piétrement, O., and Pastré, D. (2014). Free mRNA in excess upon polysome dissociation is a scaffold for protein multimerization to form stress granules. Nucleic Acids Res. 42, 8678–8691.

Burke, J.M., Moon, S.L., Matheny, T., and Parker, R. (2019). RNase L Reprograms Translation by Widespread mRNA Turnover Escaped by Antiviral mRNAs. Mol. Cell 75, 1203–1217.

Calo, E., Flynn, R.A., Martin, L., Spitale, R.C., Chang, H.Y., and Wysocka, J. (2015). Nuclear helicase DDX21 coordinates transcription and ribosomal RNA processing. Nature 518, 249–253.

Cencic, R., and Pelletier, J. (2016). Hippuristanol – a potent steroid inhibitor of eukaryotic initiation factor 4A. Translation (Austin) 4, e1137381. doi: 10.1080/21690731.2015.1137381

Chalupníková, K., Lattmann, S., Selak, N., Iwamoto, F., Fujiki, Y., and Nagamine, Y. (2008). Recruitment of the RNA helicase RHAU to stress granules via a unique RNA-binding domain. J. Biol. Chem. 283, 35186–98. b

Charroux, B., Pellizzoni, L., Perkinson, R.A., Shevchenko, A., Mann, M., and Dreyfuss, G. (1999). Gemin3: A novel DEAD box protein that interacts with SMN, the spinal muscular atrophy gene product, and is a component of gems. J. Cell Biol. 147, 1181–1194.

Chen, M.C., Tippana, R., Demeshkina, N.A., Murat, P., Balasubramanian, S., Myong, S., and Ferré-D’Amaré, A.R. (2018). Structural basis of G-quadruplex unfolding by the DEAH/RHA helicase DHX36. Nature 558, 465–469.

Delarue, M., Brittingham, G.P., Pfeffer, S., Surovtsev, I.V., Pinglay, S., Kennedy, K.J., Schaffer, M., Guiterrez, J.I., Sang, D., Poterewicz, G., Chung, J.K., Plitzko, J.M., Groves, J.T., Jacobs-Wagner, C., Engel, B.D., and Holt, L.J. (2018). mTORC1 Controls Phase Separation and the Biophysical Properties of the Cytoplasm by Tuning Crowding. Cell 174, 338–349.

Dias, A.P., Dufu, K., Lei, H., and Reed, R. (2010). A role for TREX components in the release of spliced mRNA from nuclear speckles. Nat. Commun. 1, 97 https://doi.org/10.1038/ncomms1103.

Ellis, R.J. Macromolecular crowding: obvious but unappreciated. (2001). Trends. Biochem. Sci. 26, 597–604.

Hondele, M., Sachdev, R., Heinrich, S., Wang, J., Vallotton, P., Fontoura, B.M.A., and Weis, K. (2019). DEAD-box ATPases are global regulators of phase-separated organelles. Nature 573, 144–148.

Fay, M.M., Anderson, P.J., and Ivanov, P. (2017). ALS/FTD-associated C9ORF72 repeat RNA promotes phase transitions in vitro and in cells. Cell Rep. 21, 3573–3584.

Feric, M., Vaidya, N., Harmon, T.S., Mitrea, D.M., Zhu, L., Richardson, T.M., Kriwacki, R.W., Pappu, R.V., and Brangwynne, C.P. (2016). Coexisting liquid phases underlie nucleolar subcompartments. Cell 165, 1686–1697.

Ferrandon, D., Koch, I., Westhof, E., and Nüsslein-Volhard, C. (1997). RNA-RNA interaction is required for the formation of specific bicoid mRNA 3’ UTR-STAUFEN ribonucleoprotein particles. EMBO J. 16, 1751–1758.

Fox, A.H., Lam, Y.W., Leung, A.K., Lyon, C.E., Andersen, J., Mann, M., and Lamond, A.I. (2002). Paraspeckles: a novel nuclear domain. Curr. Biol. 8, 13–25.

Frederikson, S., and Hearst, J.E. (1979). Binding of 4’-aminomethyl 4,5’,8-trimethyl psoralen to DNA, RNA, and protein in HeLa cells and *Drosophila* cells. Biochim. Biophys. Acta 563, 343–355.

Hillicker, A., Gao, Z., Jankowsky, E., and Parker, R.. The DEAD-box protein Ded1 modulates translation by the formation and resolution of an eIF4F-mRNA complex. Mol. Cell 43, 962–72.

Hochberg-Laufer, H., Schwed-Gross, A., Neuebauer, K.M., and Shav-Tal, Y. (2019). Uncoupling of nucleo-cytoplasmic RNA export and localization during stress. Nucleic Acids Res. 47, 4778–4797.

Hubstenberger, A., Noble, S.L., Cameron, C., and Evans, T.C. (2013). Translation repressors, an RNA helicase, and developmental cues control RNP phase transitions during early development. Dev. Cell 27, 161–173.

Hubstenberger, A., Courel, M., Bénard, M., Sonquere, S., Ernoult-Lange, M., Chouaib, R., Yi, Z., Morlot, J.B., Munier, A., Fradet, M., et al. (2017). P-Body Purification Reveals the Condensation of Repressed mRNA Regulons. Mol. Cell 68, 144–157.

Itzhak, D.N., Tyanova, S., Cox, J., and Borner, G.H.H. (2016). Global, quantitative and dynamic of protein subcellular localization. eLife 5, e16950.

Ivanov, P., Kedersha, N., and Anderson, P. (2019). Stress Granules and Processing Bodies in Translational Control. Cold Spring Harb. Perspect. Biol. 11, pii:a032813. doi:10.1101/cshperspect.a032813.

Jain, A., and Vale, R.D. (2017). RNA phase transitions in repeat expansion disorders. Nature 546, 243–247.

Jain, S., Wheeler, J.R., Walters, R.W., Agrawal, A., Barsic, A., and Parker, R. (2016). ATPase-modulated stress granules contain a diverse proteome and substructure. Cell 164, 487–489.

Jambor, H., Brunel, C., and Ephrussi, A. (2011). Dimerization of oskar 3’ UTRs promotes hitchhiking for RNA localization in the Drosophila oocyte. RNA 17, 2049–2057.

Jarmoskaite, I., and Russell, R. (2014). RNA helicase proteins as chaperones and remodelers. Ann. Rev. Biochem. 83, 697–725.

Kedersha, N., Cho, M.R., Lei, W., Yacono, P.W., Chen, S., Gilks, N., Golan, D.E., and Anderson, P. (2000). Dynamic shuttling of Tia-1 accompanies the recruitment of mRNA to stress granules. J. Cell Biol. 151, 1257–1268.

Kedersha, N., Panas, M.D., Achorn, C.A., Lyons, S., Tisdale, S., Hickman, T., Thomas, M., Lieberman, J., McInnery, G.M., Ivanov, P., and Anderson, P. (2016). G3BP-Caprin1-USP10 complexes mediate stress granule condensation and associate with 40S subunits. J. Cell Biol. 212, 845–60.

Kedersha, N., Stoecklin, G., Ayodele, M., Yacono, P., Lykke-Andersen, J., Fritzler, M.J., Scheuner, D., Kaufman, R.J., Golan, D.E., and Anderson, P. (2005). Stress granules and processing bodies are dynamically linked sites of mRNP remodeling. J. Cell Biol. 169, 871–84.

Khong, A., Jain, S., Matheny, T., Wheeler, J.R., and Parker, R. (2017b). Isolation of mammalian stress granule cores for RNA-seq analysis. Methods 137, 49–54.

Khong, A., Matheny, T., Jain, S., Mitchell, S.F., Wheeler, J.R., and Parker, R. (2017a). The Stress Granule Transcriptome Reveals Principles of mRNA Accumulation in Stress Granules. Mol. Cell 68, 808–820.

Khong, A., and Parker, R. (2018). mRNP architecture in translating and stress conditions reveals an ordered pathway of mRNP compaction. J. Cell Biol. 217, 4124–4140.

Langdon, E.M., Qiu, Y., Ghanbari Niaki, A., McLaughlin, G.A., Weidmann, C.A., Gerbich, T.M., Smith, J.A., Crutchley, J.M., Termini, C.M., Weeks, K.M., et al. (2018). mRNA structure determines specificity of a polyQ-driven phase separation. Science 360, 922–927.

Linder, P., and Fuller-Pace, F.V. (2013). Looking back on the birth of DEAD-box RNA helicases. Biochim. Biophys. Acta 1829, 750–5.

Lindqvist, L., Oberer, M., Reibarkh, M., Cencic, R., Bordeleau, M.E., Vogt, E., Marintchev, A., Tanaka, J., Fagotto, F., Altmann, M., Wagner, G., and Pelletier, J. (2008). Selective pharmacological targeting of a DEAD-box helicase. PLoS One 3, e1583.

Liu, F., Putnam, A., and Jankowsky, E. (2008). ATP hydrolysis is required for DEAD-box protein recycling but not for unwinding. Proc. Natl. Acad. Sci. USA 105, 20209–20214.

Lorsch, J.R., and Herschlag, D. (1998a). The DEAD box protein eIF4A. 1. A minimal kinetic and thermodynamic framework reveals coupled binding of RNA and nucleotide. Biochemistry 37, 2180–2193.

Lorsch, J.R., and Herschlag, D. (1998b). The DEAD box protein eIF4A. 2. A cycle of nucleotide and RNA-dependent conformational changes. Biochemistry 37, 2194–2206.

Markmiller, S., Soltanieh, S., Server, K.L., Mak, R., Jin, W., Fang, M.Y., Luo, E.C., Krach, F., Yang, D., Sen, A., et al. (2018). Context-dependent and disease-specific diversity in protein interactions within stress granules. Cell 172, 590–604.

Mahadevan, K., Zhang, H., Akef, A., Cui, X.A., Gueroussov, S., Cenik, C., Roth, F.P., and Palazzo, A.F. (2013). RanBP2/Nup358 potentiates the translation of a subset of mRNAs encoding secretory proteins. PLoS Biol. 11, e1001545 doi:10.1371/journal.pbio.1001545

Mogk, A., Bukau, B., and Kampinga, H.H. (2018). Cellular handling of protein aggregates by disaggregation machines. Mol. Cell 69, 214–226.

Moon, S.L., Morisaki, T., Khong, A., Lyon, K., Parker, R., and Stasevich, T.J. (2019). Multicolour single-molecule tracking of mRNA interactions with RNP granules. Nat. Cell Biol. 21, 162–168.

Murat, P., Marisco, G., Herdy, B., Ghanbarian, A.T., Portella, G., and Balasubramanian, S. (2018). RNA G-quadruplexes at upstream open reading frames cause DHX36- and DHX9-dependent translation of human mRNAs. Genome Biol. 19, 229.

Oguro, A., Ohtsu, T., Svitkin, Y.V., Sonenberg, N., and Nakamura, Y. (2003). RNA aptamers to initiation factor 4A helicase hinder cap-dependent translation by blocking ATP hydrolysis. RNA 9, 394–407.

Pause, A., Méthot, N., Svitkin, Y., Merrick, W.C., and Sonenberg, N. (1994). Dominant negative mutants of mammalian translation initiation factor eIF-4A define a critical role for eIF-4F in cap-dependent and cap-independent initiation of translation. EMBO J. 13, 1205–1215.

Protter, D.S.W., and Parker, R. (2016). Principles and Properties of Stress Granules. Trends Cell Biol. 26, 668–679.

Putnam, A.A., and Jankowsky, E. (2013). DEAD-box helicases as integrators of RNA, nucleotide, and protein binding. Biochim. Biophys. Acta 1829, 884–893.

Ribero de Almeida, C., Dhir, S., Dhir, A., Moghaddam, A.E., Sattentau, Q., Meinhart, A., Proudfoot, N.J. (2018). RNA Helicase DDX1 Converts RNA G-Quadruplex Structures into R-Loops to Promote IgH Class Switch Recombination. Mol. Cell 70, 650–662.

Rogers, G.W., Jr., Richter, N.J., and Merrick, W.C. (1999). Biochemical and kinetic characterization of the RNA helicase activity of eukaryotic initiation factor 4A. J. Biol. Chem 274, 122236–12244.

Rogers, G.W., Jr., Lima, W.F., and Merrick, W.C. (2001). Further characterization of the helicase activity of eIF4A. Substrate specificity. J. Biol. Chem. 276, 12598–12608.

Rowlinson, J.S., and Widom, B. (1982). Molecular Theory of Capillarity. (New York: Oxford University Press).

Rueden, C.T., Schindelin, J., Hiner, M.C., DeZonia, B.E., Walter, A.E, and Eliceiri, K.W. (2017). ImageJ2: ImageJ for the next generation of scientific data. BMC Bioinformatics 18, 529.

Saitoh, N., Spahr, C.S., Patterson, S.D., Bubulya, P., Neuwald, A.F., and Spector, D.L. (2004). Proteomic analysis of interchromatin granule clusters. Mol. Biol. Cell 15, 3876–3890.

Sauer, M., Juranek, S.A., Marks, J., De Magis, A., Kazemeier, H.G., Hilbig, D., Benhalevy, D., Wang, X, Hafner, M., and Paeschke, K. (2019). DHX36 prevents the accumulation of translationally inactive mRNAs with G4-structures in untranslated regions. Nat. Commun. 10, 2421. doi:10.1038/s41467-019-10432-5

Schindelin, J., Arganda-Carreras, I., Frise, E., Kaynig, V., Longair, M., Pietzsch, T., Rueden, C., Saalfeld, S., Schmid, B., et al. (2012). Fiji: an open-source platform for biological-image analysis. Nat. Methods 9, 676–682.

Shapiro, E., Biezuner, T., and Linnarsson, S. (2013). Single-cell sequencing-based technologies will revolutionize whole-organism science. Nat. Rev. Genet. 14, 618–630.

Shin, Y., and Brangwynne, C.P. (2017). Liquid phase condensation in cell physiology and disease. Science 357, eaaf4382 doi: 10.1126/science.aaf4382.

Singh, A.K., Choudhury, S.R., De, S., Zhang, J., Kissane, S., Dwivedi, V., Ramanthan, P., Petric, M., Orsini, L., Hebenstreit, D., and Brogna, S. (2019). The RNA helicase UPF1 associates with mRNAs co-transcriptionally and is required for the release of mRNAs from gene loci. eLife 8, e41444.

Sun, Y., Atas, E., Lindqvist, L.M., Sonenberg, N., Pelletier, J., and Meller, A. (2014). Single-molecule kinetics of the eukaryotic initiation factor 4AI upon RNA unwinding. Structure 22, 941–948.

Svitkin, Y.V., Pause, A., Haghighat, A., Pyronnet, S., Witherell, G., Belsham, G.J., and Sonenberg, N. (2001). The requirement for eukaryotic initiation factor 4A (eIF4A) in translation is in direct proportion to the degree of mRNA 5’ secondary structure. RNA 7, 382–394.

Tanaka, T., Ohashi, S., and Kobayashi, S. (2014). Roles of YB-1 under arsenite-induced stress: translational activation of HSP70 mRNA and control of the number of stress granules. Biochim. Biophys. Acta 1840, 985–992.

Tauber, D., and Parker, R. (2019). 15-Deoxy-Δ^12,14^-prostaglandin J2 promotes phosphorylation of eukaryotic initiation factor 2α and activates the integrated stress response. J. Biol. Chem. 294, 6344–6352.

Tourrière, H., Chebli, K., Zekri, L., Courselaud, B., Blanchard, J.M., Bertrand, E., and Tazi, J. (2003). The RasGAP-associated endoribonuclease G3BP assembles stress granules. J. Cell Biol. 160, 823–831.

Trcek, T., Grosch, M., York, A., Schroff, H., Lionnet, T., and Lehmann, R. (2015). Drosophila germ granules are structured and contain homotypic mRNA clusters. Nat. Commun. 6, 7962.

Tu, Y.T., & Barrientos, A. (2015). The Human Mitochondrial DEAD-Box Protein DDX28 Resides in RNA Granules and Functions in Mitoribosome Assembly. Cell Rep. 10, 854–864.

Van Treeck, B., and Parker, R. (2018). Emerging Roles for Intermolecular RNA-RNA Interactions in RNP Assemblies. Cell 174, 791–802.

Van Treeck, B., Protter, D.S.W., Matheny, T., Khong, A., Link, C.D., and Parker, R. (2018). RNA self-assembly contributes to stress granule formation and defining the stress granule transcriptome. Proc. Natl. Acad. Sci. USA 115, 2734–2739.

West, J.A., Mito, M., Kurosaka, S., Takumi, T., Tanegashima, C., Chujo, T., Yanaka, K., Kingston, R.E., Hirose, T., Bond, C., et al. (2016). Structural, super-resolution microscopy analysis of paraspeckle nuclear body organization. J. Cell Biol. 214, 817–30.

Youn, J.Y., Dunham, W.H., Hong, S.J., Knight, J.D.R., Bashkurov, M., Chen, G.I., Bagci, H., Rathod, B., MacLeod, G., Eng, S.W.M., et al. (2018). High-density proximity mapping reveals the subcellular organization of mRNA-associated granules and bodies. Mol. Cell 69, 517–532.

Zhang, Y., Burkhardt, D.H., Rouskin, S., Li, G.W., Weissman, J.S., and Gross, C.A. (2018). A Stress Response that Monitors and Regulates mRNA Structure is Central to Cold Shock Adaptation. Mol. Cell 70, 274–286.

Zimmerman, S.B. (1993). Macromolecular crowding effects on macromolecular interactions: some implications for genome structure and function. Biochim. Biophys. Acta 1216, 175–185.

