## Supplementary figures and images for "Modulation of RNA condensation by the DEAD-box protein eIF4A"

### Supplemental Figure 1

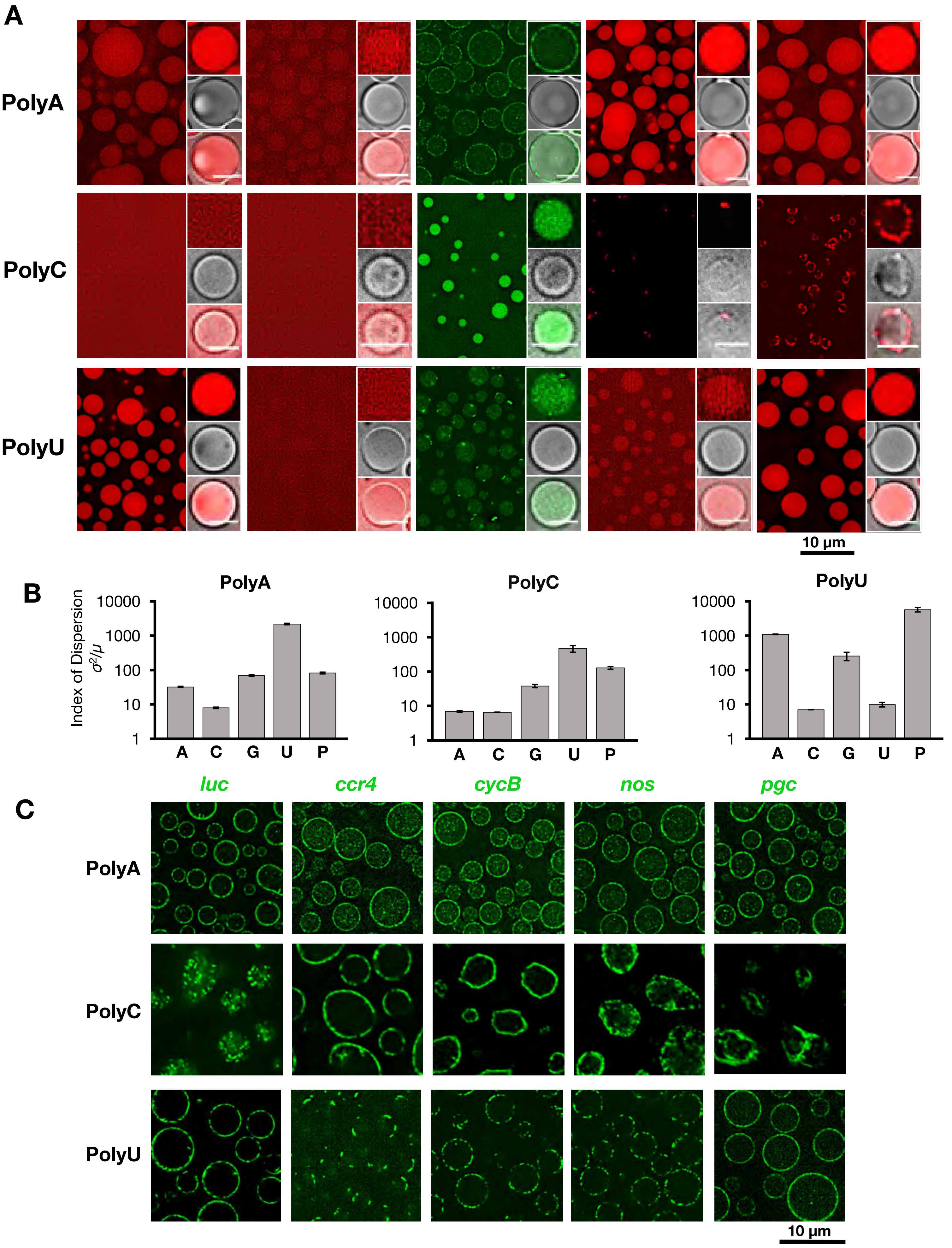

### Supplemental Figure 2

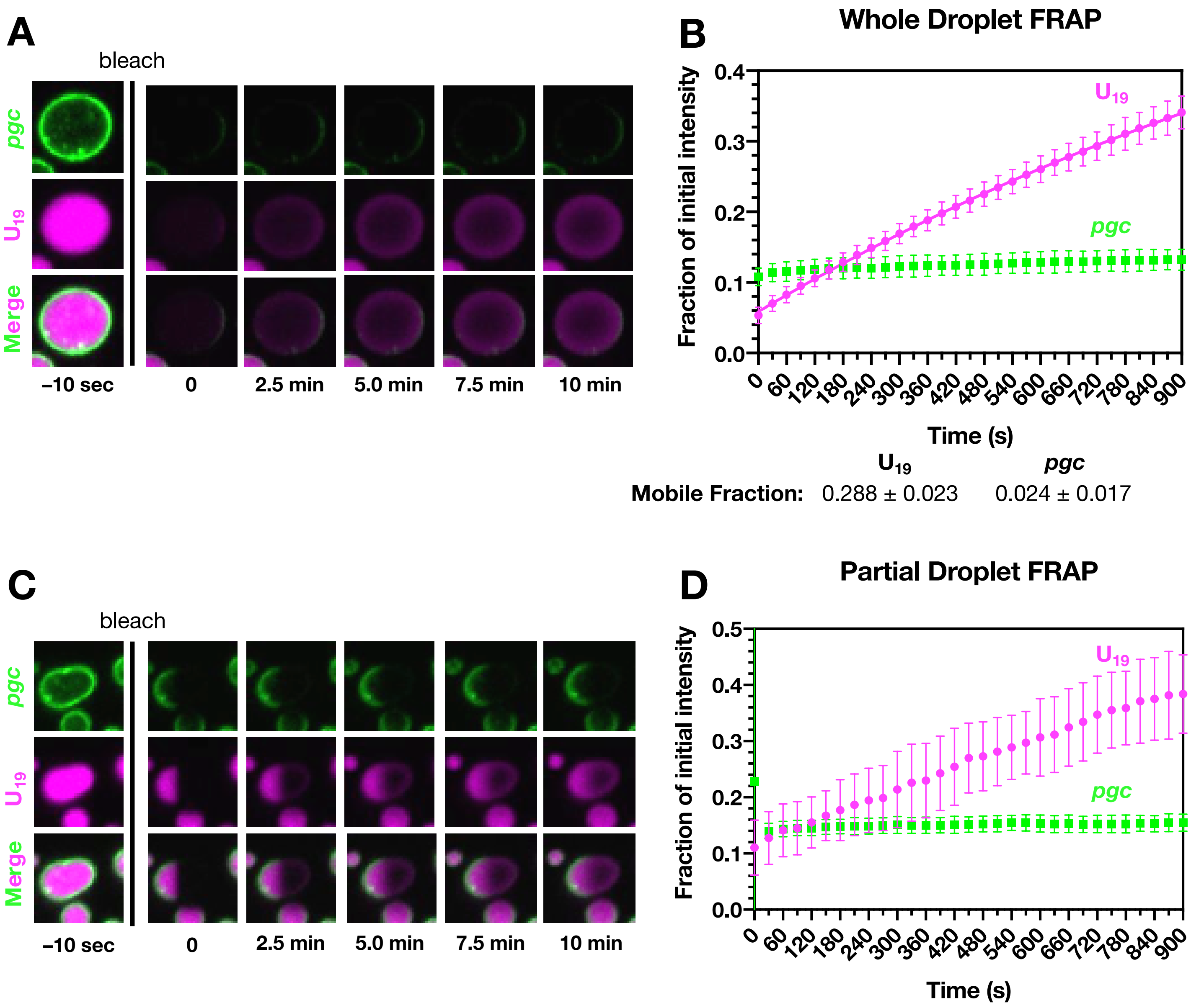

### Supplemental Figure 3

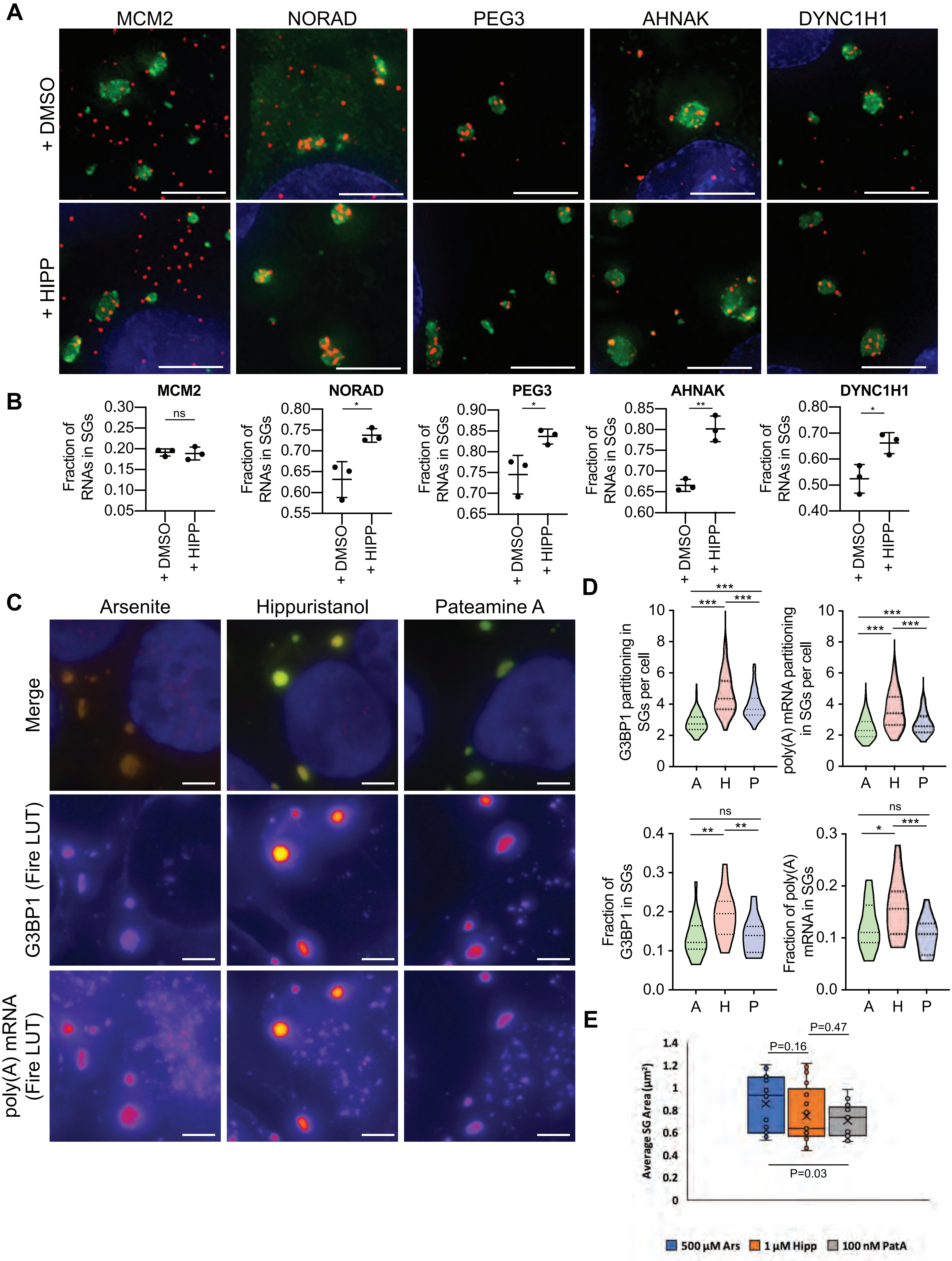

### Supplemental Figure 4

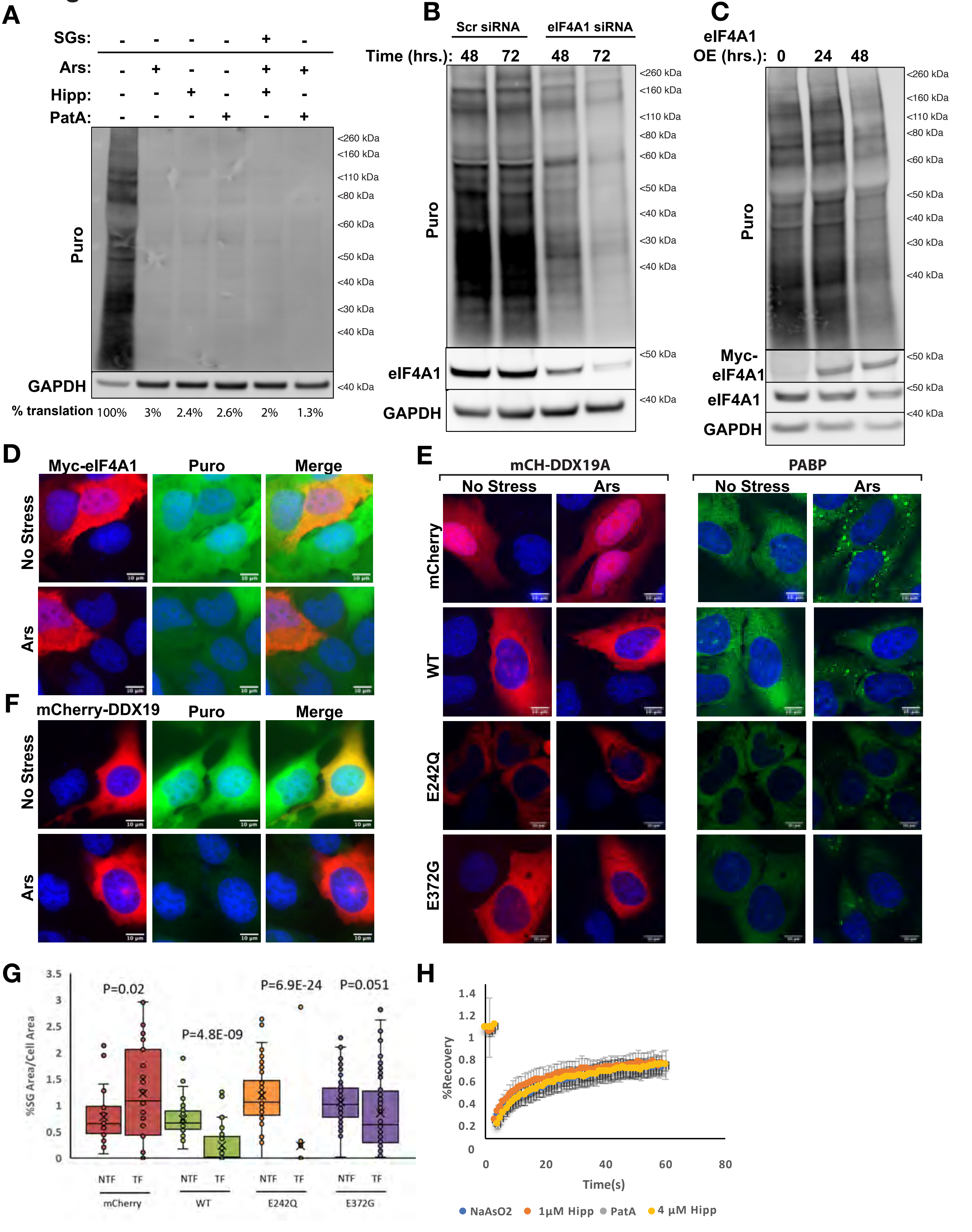

### Supplemental Figure 5

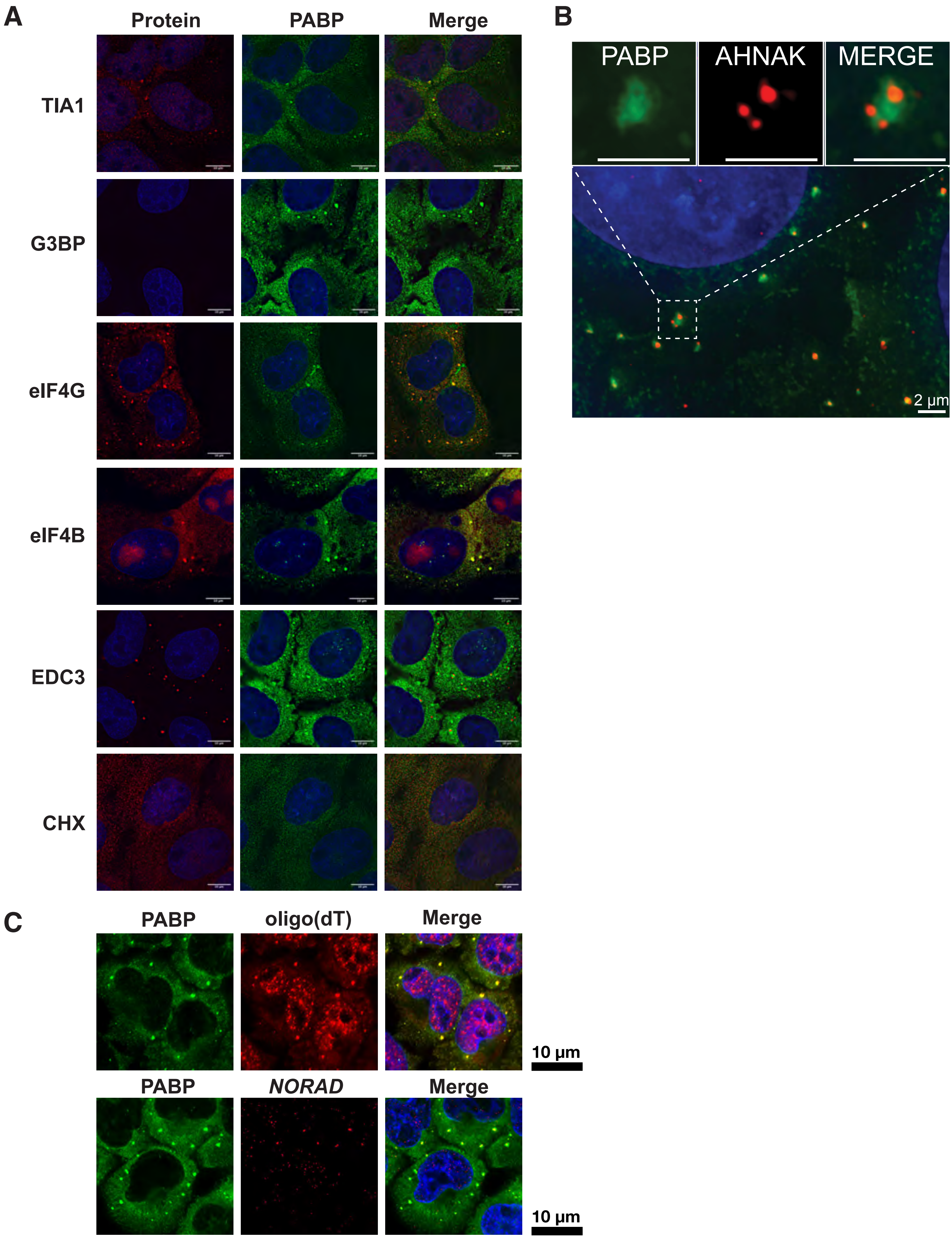

### Supplemental Figure 6

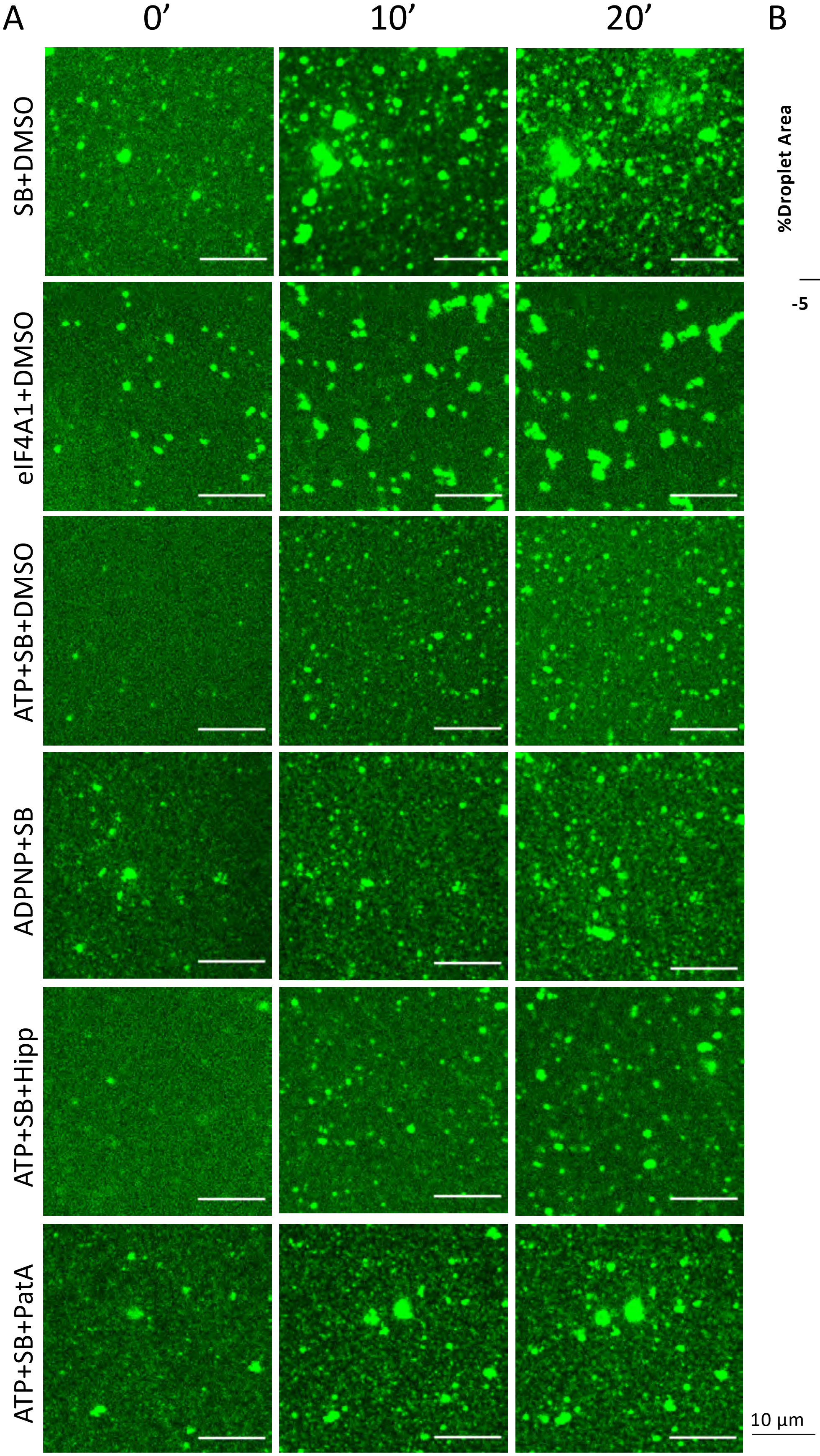

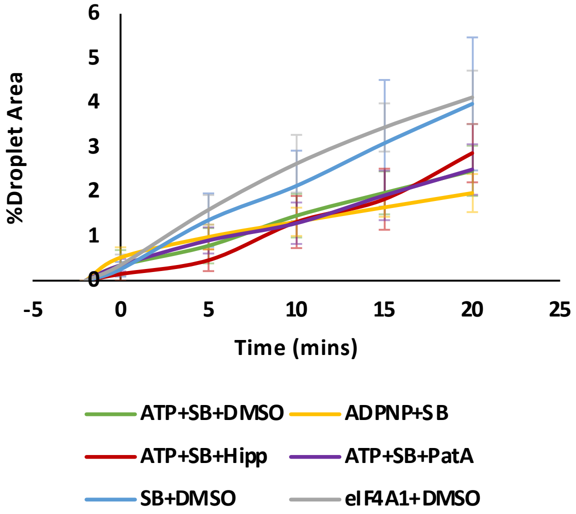
